# Sleep specific changes in infra-slow and respiratory frequency drivers of cortical EEG rhythms

**DOI:** 10.1101/2023.01.20.524831

**Authors:** Tommi Väyrynen, Heta Helakari, Vesa Korhonen, Johanna Tuunanen, Niko Huotari, Johanna Piispala, Mika Kallio, Lauri Raitamaa, Janne Kananen, Matti Järvelä, J. Matias Palva, Vesa Kiviniemi

## Abstract

Infra-slow fluctuations (ISFs, 0.008-0.1 Hz) characterize hemodynamic and electric potential signals from the human brain. ISFs are known to correlate with the amplitude dynamics of fast (> 1 Hz) neuronal oscillations, and may arise from permeability fluctuations of the blood-brain barrier (BBB). Slow physiological pulsations such as respiration may also influence the amplitude dynamics of fast oscillations, but it remains uncertain if these processes track the fluctuations of fast cortical oscillations or act as their drivers. Moreover, possible effects of sleep and associated BBB permeability changes on such coupling are unknown. Here, we used non-invasive high-density full-band electroencephalography (EEG) in healthy human volunteers (N=21) to measure concurrently the ISFs, respiratory pulsations, and fast neuronal oscillations during periods of wakefulness and sleep, and to assess the strength and direction of their phase-amplitude coupling. The phases of ISFs and respiration were both coupled with the amplitude of fast neuronal oscillations, with stronger ISF coupling evident during sleep. Causality analysis robustly showed that the phase of ISF and respiration drove the amplitude dynamics of fast oscillations in sleeping and waking states. However, the net direction of modulation was stronger during the awake state, despite the stronger power and phase-amplitude coupling of slow signals during sleep. These findings show that the ISFs in slow cortical potentials and respiration together significantly determine the dynamics of fast cortical oscillations. We propose that these slow physiological phases are involved in coordinating cortical excitability, which is a fundamental aspect of brain function.

**Significance Statement:** Previously disregarded EEG infra-slow fluctuations (0.008-0.1 Hz) and slow physiological pulsations such as respiration have been attracting increasing research interest, which shows that both of these signals correlate with fast (> 1 Hz) neuronal oscillations. However, little has been known about the mechanisms underlying these interactions; for example, the direction of causality in this interaction has not hitherto been studied. Therefore, we investigated full-band EEG in healthy volunteers during wakefulness and sleep to determine if ISF and respiration phases drive neuronal amplitudes. Results showed that ISF and respiration are phase-amplitude coupled, and predict neuronal EEG rhythms. Thus, we conclude that fast neuronal rhythms in human brain are modulated by slower non-neural phenomena.

## Main Text

Quasiperiodic infra-slow fluctuations (ISFs, 0.008-0.1 Hz) (Vanhatalo et al., 2004; Hughes et al., 2011; Palva and Palva, 2012), comprise the strongest signal in full-band electroencephalography (fbEEG) (Niedermeyer et al., 2011). ISFs arise largely from non-neuronal sources such as alterations in cerebral blood flow (Tschirgi and Taylor, 1958; Held et al., 1964; Besson et al., 1970). Results of studies in animal models (Nita et al., 2004) and humans (Kiviniemi et al., 2017) suggest that the dynamics of blood-brain barrier (BBB) permeability could be a major contributor to the EEG ISF signal. In awake state, the fbEEG ISFs are correlated with statistically independent ISFs in connected networks of the blood-oxygenation-level dependent (BOLD) signal during functional magnetic resonance imaging (fMRI), (Leopold et al., 2003; Hiltunen et al., 2014).

In addition to ISF, there is also a link between respiration and fast neuronal changes as measured with intracranial needle electrode recordings (Iwabe et al., 2014; Zelano et al., 2016; Herrero et al., 2018), motor activity recordings (Cao et al., 2012) and magnetoencephalography (MEG) measurements (Kluger and Gross, 2021). In mice, the respiratory cycle correlates with cortical neuronal activity (Ito et al., 2014), excitability (Li and Rymer, 2011), and sensory perception (Flexman et al., 1974). The power in ISF and respiratory frequency BOLD signal pulsations increases during sleep in brain areas anatomically overlapping with regions showing concomitant EEG slow-wave increases (Helakari et al., 2022).

Infraslow 0.034 Hz oscillations of norepinephrine release from locus coeruleus orchestrates sleep architecture by modulating neuronal activity across a wide dynamic range (Osorio-Forero et al., 2021; Kjaerby et al., 2022). Interestingly, both the magnitude of ISFs in the EEG (Marshall et al., 1996, 1998) and neurovascular BOLD signal (Fukunaga et al., 2006; Fultz et al., 2019) are greater during sleep than in the awake state. This is in line with the hypothesis that circadian fluctuations in BBB function contribute to ISF-signal generation, as seen in studies with sleep deprivation (Cuddapah et al., 2019). Moreover, experimentally increasing the flux of cerebrospinal fluid (CSF), which is associated with elevated BBB permeability in mice, can shift the EEG from the awake activity pattern into a sleep-like state, which is dominated by slow-wave activity (Ding et al., 2016). It has also been shown that the BBB permeability increases as a function of isoflurane anesthesia depth, reflected as an increase in EEG ISFs (Tétrault et al., 2008).

Thus, several lines of evidence concur in linking slow physiological signals in fbEEG with fast (> 1 Hz) neuronal activities during sleep (Vanhatalo et al., 2004) and awake (Monto et al., 2008) states, such that the phase of ISFs correlates with the amplitude of fast neuronal rhythms. Furthermore, ISF and respiratory phase also correlated with behavioral performance in a stimulus-detection task with humans (Monto et al., 2008; Johannknecht and Kayser, 2022), which implies that the EEG derives functionally significant contributions from these slow, non-neuronal physiological signals. However, the directionality of causality in the relationship between the slow physiological signals and fast neuronal oscillations remains unknown in sleep and awake brain states.

Here, our first objective was to assess how the frequency characteristics of the EEG changes in sleep. Second, we asked whether the ISFs (ISF_EEG_) and respiration (RESP_EEG_) induced EEG signals arise as a “byproduct” of emergent slow dynamics (Palva et al., 2013) of fast neuronal oscillations, or rather play a causal role in modulating the fast neuronal oscillations *per se*. To further elucidate the underlying mechanistic basis of these interactions, our final objective was to uncover whether coupling between fast neuronal processing and non-neuronal slow processes would be altered during non-rapid eye movement (NREM) sleep. To test these conjectures, we assessed spectral power, phase-amplitude coupling, and directional drive between ISF_EEG_ and fast oscillations in human brain using 256-channel fbEEG during wakefulness and sleep. The results show that slow physiological oscillation phases both couple and drive fast neurophysiological rhythm amplitudes across awake and sleep states in healthy human volunteers.

## Results

### Infra-slow frequency power increases during sleep

Our hypothesis was that EEG low frequency power should increase in sleep in concert with known increases in ISF BOLD signal power. To determine whether the frequency characteristics of the signal change in NREM sleep, we used spectral analysis to assess the differences in the power in infra-slow and higher frequency bands between sleep and awake states (Figure 1a). During sleep, whole head spectral power was higher in the ISF_EEG_ and RESP_EEG_ frequencies between 0.02-0.25 Hz (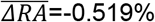, *p*<0.05, *η*^2^=0.331), and conversely was lower in frequencies above 1.9 Hz (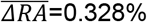, *p*<0.05, *η*^2^=0.309). Respiratory pulsation peaks occurred at ~0.3 Hz in the awake brain versus ~0.2 Hz during sleep. Slow-wave (0.2-2 Hz) EEG topography during sleep showed increased power extending over most of the cortical surface, excluding frontal electrodes. In line with our hypothesis, ISF power was increased over wide areas in sleep, with pronounced increases in frontal regions. In contrast, the high frequency power dropped, suggesting a shift of the power of cortical oscillations from high to low frequencies in NREM sleep.

**Figure 1.**
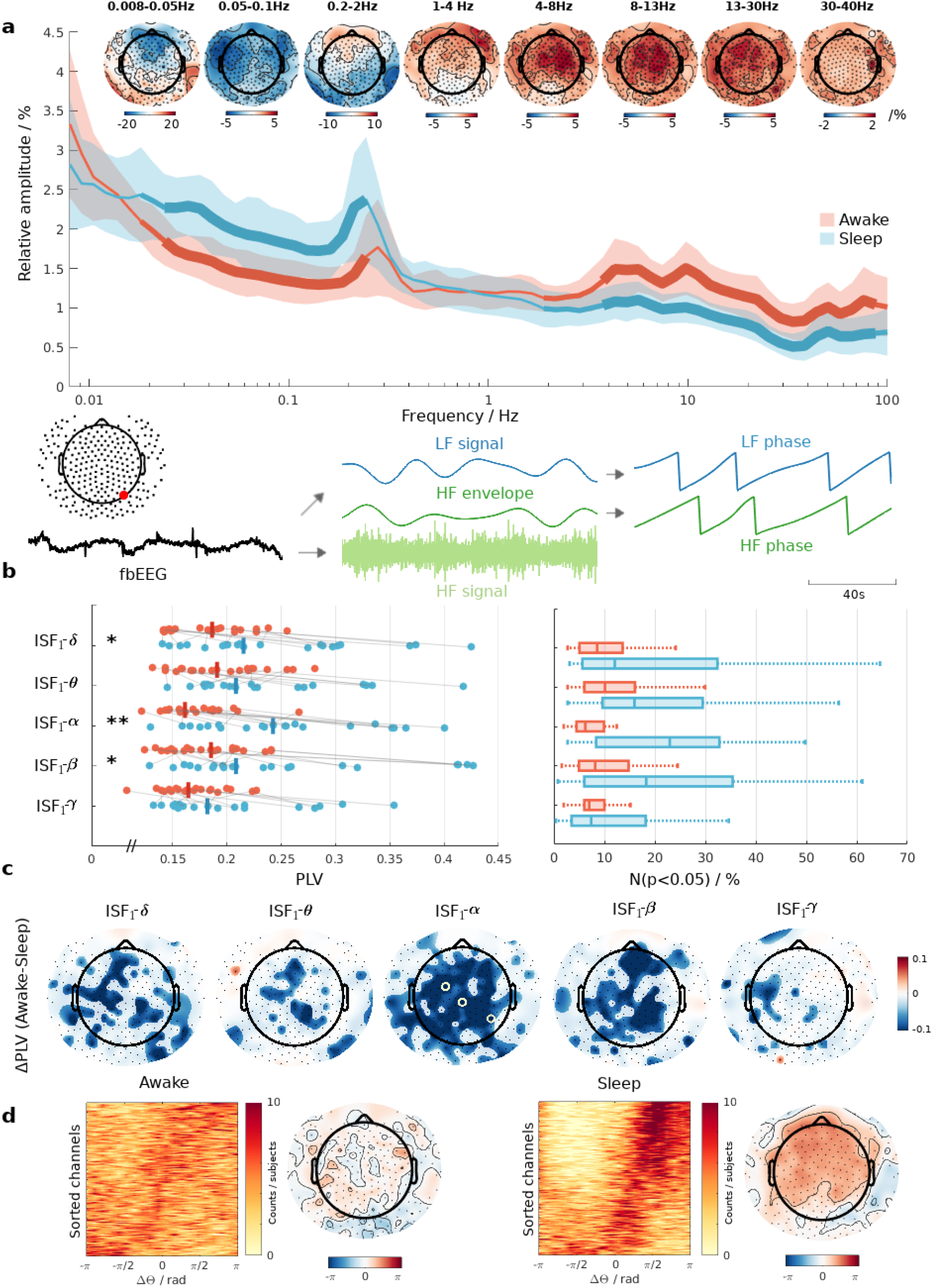
Effect of sleep on oscillation power and phase-amplitude coupling with cortical rhythms. a) EEG relative amplitude on a logarithmic frequency scale, where shadings around the mean are standard deviations of the groups. The width of the solid lines represents statistical significance (p<0.05) in rising order of rank: no statistical significance, permutation tested, permutation test + maximum statistics correction. Topography plots the relative power difference (awake-sleep) at ISF (<0.01 Hz), slow-wave (0.2-2 Hz) and neural frequency bands. Bottom: Illustration of phase-amplitude coupling with one EEG trace. b) Left: Median phase-locking values (PLV) taken over electrodes. Asterisks indicate statistically significant (adjusted p<0.05*, 0.01**, 0.001***) difference in the coupling strength between the two states. Gray lines connect paired subjects. Right: Number of significantly coupled electrodes for each subject, scaled to percentiles. c) Difference in average PLV combined with permutation testing (p<0.05) whitening mask. Yellow circles around electrodes indicate significance with maximum statistic correction (p<0.05). d) Probability estimates of the average phase difference between infra-slow fluctuations (ISF) phase and fast rhythms phase combined from all fast frequency bands. Channels on the y-axis are sorted to ascending order following the median phase difference. Topography plots the median phase difference taken over the fast bands and subjects.

### Sleep increases ISF coupling with cortical rhythms

We next supposed that the coupling between the physiological slow brain fluctuations and neuronal oscillations would increase in sleep. To assess this relationship between the ISF phase and the amplitudes of fast oscillations, we employed a phase-amplitude coupling estimator based on phase-locking value (PLV) (Vanhatalo et al., 2004), which enables the isolation of phase-amplitude coupling effects attributable to distinct ISF frequencies in the amplitude time series. Since phase locking is a property of narrow-band signals, we split the ISF_EEG_ into two separate frequency bands (ISF_1-EEG_: 0.008-0.05 Hz, and ISF_2-EEG_: 0.05-0.1 Hz) to improve our analysis accuracy. We present the ISF_2-EEG_ related results in the Supplementary section (Figures S2 & S3).

In line with our hypothesis, the phase-amplitude coupling between the phase of ISF_1-EEG_ and the amplitude of all fast rhythms was greater during sleep than during the awake recordings (Figure 1b, left). This difference as a function of arousal state was strongest in the alpha band. Groupwise difference in phase-amplitude coupling was significant between the ISF_1-EEG_ phase and the amplitude of delta (*ΔPLV*=-0.029, *p_adj_*=0.031, *η^2^*=0.179), alpha (*ΔPLV*=-0.081, *p_adj_*=0.002, *η^2^*=0.339), and beta (*ΔPLV*=-0.023, *p_adj_*=0.037, *η^2^*=0.142) bands, which were focused on central brain regions (Figure 1c). Theta and gamma bands showed similar trends, with stronger phase locking during sleep. PLV compared with surrogate data followed the same pattern as did raw PLVs (Figure 1b, right). The interquartile range for the number of significantly coupled scalp electrodes was clearly lower during wakefulness, when only 5-15 % of electrodes had significant phase-amplitude coupling effects as compared to 5-35 % during sleep.

To investigate the coherence of the phase differences, we estimated the probability distributions of the average phase-amplitude coupling phase differences combined from all frequency bands (Figure 1d & S6) between the ISF_1-EEG_ and fast oscillations. During wakefulness, the phase differences were distributed in a relatively uniform manner. However, the pattern changed drastically during sleep, when there was a distinct tendency towards phase differences around π2, which corresponded to the falling phase of the ISF cycle. Topographical analysis showed that these changes were focused on parietal, central and frontal electrodes.

Taken together, these results show that, while ISFs are significantly coupled to the amplitude dynamics of fast neuronal oscillations during both awake and sleep states, this relationship is stronger and more widespread across the cortical surface during sleep.

### ISF phase drives electrophysiological brain rhythms

We next asked whether there was any directed net drive, such that ISFs were driving the cortical amplitude fluctuations, rather than emerging as their consequence. To resolve this question, we calculated directed phase-amplitude coupling according to phase transfer entropy (PTE) between the phase of ISFs and the neural amplitudes. Upon assessing PTE in both possible directions of interaction (Figure 2a), we found increased correlations for both directions and in all frequency bands during the awake state in comparison to sleep.

**Figure 2.**
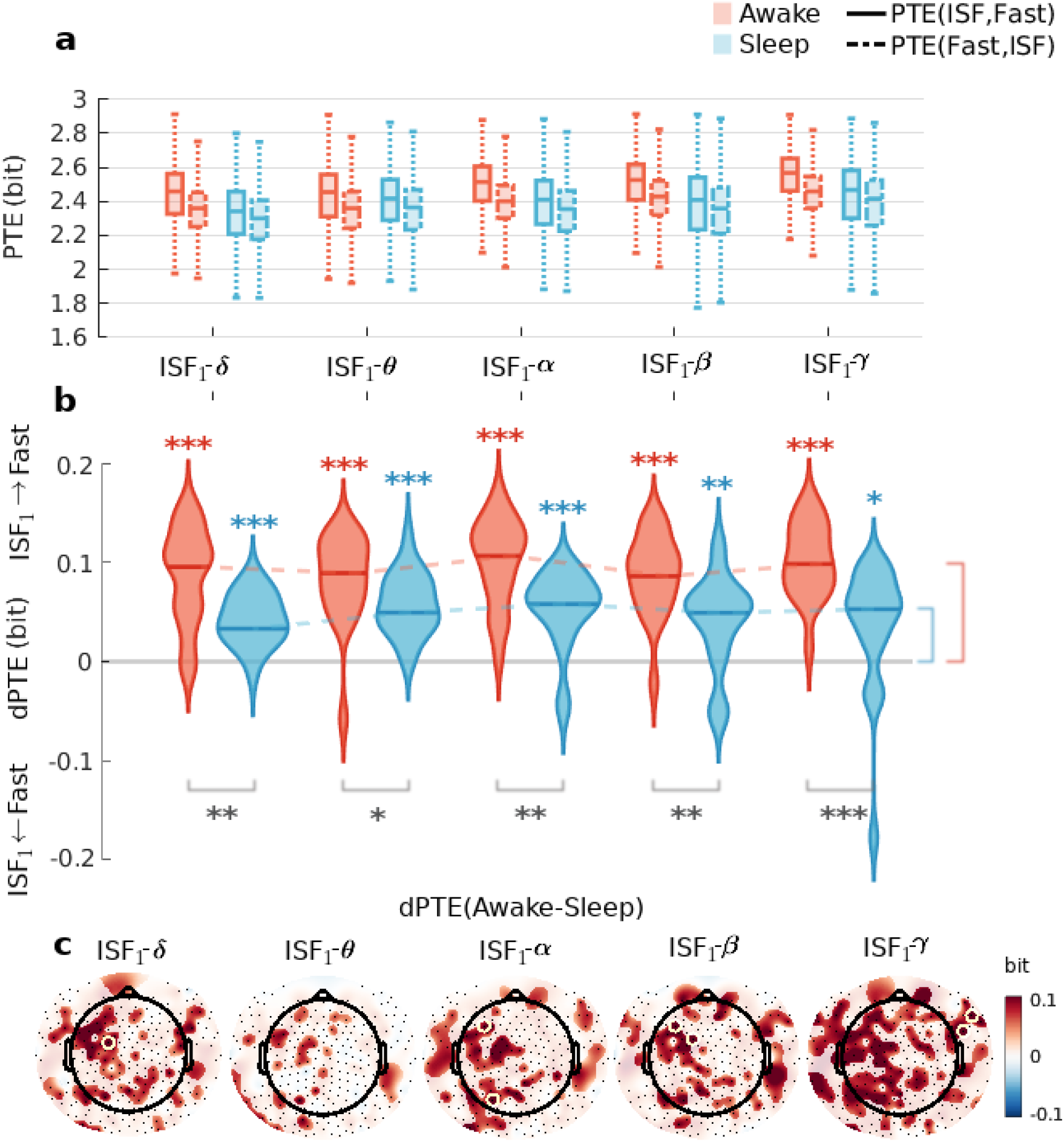
The phase of ISF1 drives infra-slow amplitude fluctuations of fast cortical oscillations. a) Box plot of phase transfer entropy (PTE) for both directions: ISF-neural & neural-ISF, where the box contains the interquartile range (IQR) and whiskers 1.5*IQR. b) Probability density estimate of the average directional PTE (dPTE). Since dPTE is a directional metric, the net prediction direction changes on a sign change. Greek letters indicate the cortical band in question. Asterisks indicate statistical significance: colored asterisks are for one-sample tests marking significant non-zero drive. Black asterisks are for two-sample groupwise comparison. c) Topography shows average dPTE difference (awake-sleep) combined with overlayed mask, highlighting significance for permutation testing (p<0.05). Significant electrodes for maximum statistics correction (p<0.05) are visualized by yellow circles.

To identify whether the slow physiological or fast neuronal oscillations were the driver of the other, we configured PTE into a directional form. This approach revealed significant directional PTE (dPTE) in all frequency bands during waking state (combined *dPTE*: 0.099 bits, *p_adj_*<0.001 for all bands) and likewise in sleep (combined *dPTE*: 0.052 bits, *p_adj_*<0.001 for all, except gamma and beta) states (Figure 2b). Importantly, the phase of ISF was a stronger predictor of the fast neuronal rhythms than vice versa, which implies that ISFs do indeed drive the amplitude dynamics of fast oscillations. Median dPTE was consistently stronger during wakefulness than during sleep

(average difference 0.047 bits, *p_adj_*<0.02 and *η^2^* >0.16 for all bands), with little variation in the magnitude of dPTE between frequency bands.

There was decreased net drive during sleep, which arose from a drop in the correlation between ISF with neural direction, and a simultaneous smaller decreased correlation in the opposite interaction direction (Figure 2a). Scalp topography of the groupwise difference in dPTE (Figure 2c) highlights the involvement of spatially overlapping regions, which was least evident in the theta frequency band. These findings constitute the first evidence that ISFs drive cortical brain rhythms during both wakefulness and sleep states.

In general, increased coupling should reflect increased information transfer between coupled oscillators (Ceguerra et al., 2011). We initially thought that the stronger phase-amplitude coupling observed in sleep must have lowered directional drive due to more constant phase difference between the phase time series. To test this conjecture, we quantified the relationship between phase locking strength and prediction magnitudes. This analysis revealed no linear correlation between the two factors (Figure S5).

We next asked whether the spectral power increase of ISF during sleep pointed to elevated autocorrelations, and might thereby underlie the changes in directional drive. However, the autocorrelations in terms of transfer entropy did not correlate with power, and thus did not explain the difference in dPTE between arousal states (Figure S4), leading us to reject that hypothesis. Prior studies with Kuramoto models have shown that information transfer can drop significantly even before coupled oscillators have reached a full synchronization (Ceguerra et al., 2011). Thus, the reduced information transfer we observed during sleep could indicate an approach towards such a stable state.

Another theory to explain the reduced net drive of the neuronal amplitudes in sleep invokes the proposal that the interaction becomes more bidirectional in sleep, in conjunction with an increased water permeability of the BBB (and consequently reduced neural excitability), which would tend to cancel out the net drive. Nonetheless, the prediction decreased in both directions (PTE, fast→slow and slow→fast) during sleep, along with a reduced net drive as indicated by dPTE, together suggesting that the interaction does not become more bidirectional as compared to the awake state.

### EEG respiratory phase is a driver for fast oscillation amplitudes

We have recently found that respiratory brain pulsations increase in the magnetic resonance encephalography (MREG) BOLD signal and overlap with fronto-parietal slow-wave electrophysiological changes in sleep (Helakari et al., 2022). That result, along with the evident power peaks at respiratory frequencies (Figure 1a), led us to ask whether the respiratory rhythm RESP_EEG_ also exhibits coupling with fast cortical amplitude oscillations, and if sleep has any effect on this association. To address this, we repeated same analysis as above with ISF to evaluate phase-amplitude coupling using the individual respiration frequency EEG.

We found respiration frequencies to be higher during wakefulness compared to sleep (Figure 3c). The mean level of phase-amplitude coupling was stronger during wakefulness (Figure 3a), but the differences in PLV magnitudes were small, without showing significant differences between arousal states. PLV compared against surrogate data revealed that RESP_EEG_ coupling to neural amplitudes was a widespread phenomenon. Awake state recordings showed an interquartile range of significantly coupled channels ranging from 15% to as high as 80%. In contrast, the same metric ranged from 15 to only 60% during sleep. Even though whole head differences were non-significant, significant PLV differences for individual electrodes were found with the slowest delta band in frontal regions, and for theta frequencies in occipital areas (Figure 3b).

**Figure 3.**
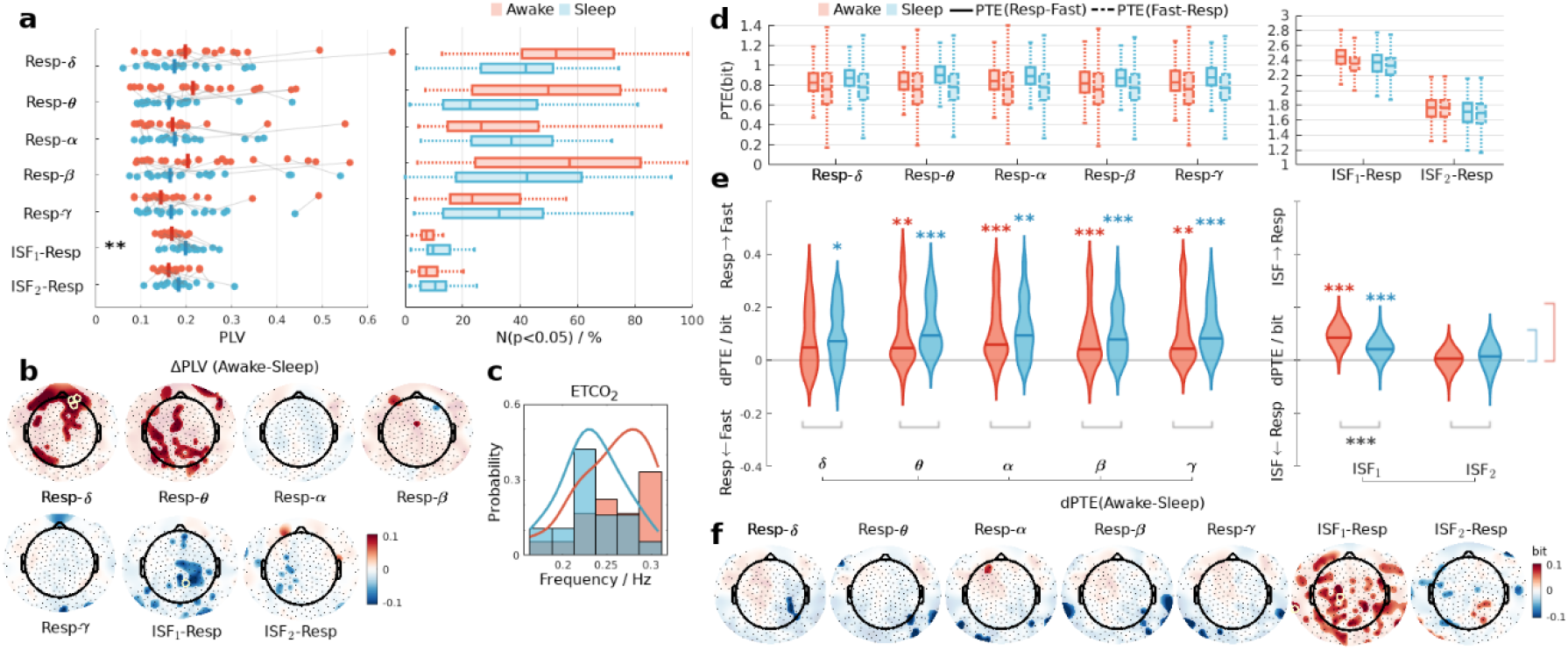
EEG respiratory frequency coupling and drive. a) On the left, the scatter plot of phase-amplitude coupling showing median phase-locking values (PLV) for each subject and arousal states. Asterisks mark significance for two-sample tests. On the right, the number of significantly coupled electrodes, scaled to percentiles. b) Difference in average PLV combined with permutation testing (p<0.05) whitening mask. Yellow circles around electrodes indicate significance with maximum statistical correction (p<0.05). c) Probability estimates of the individual respiratory frequencies taken from end-tidal CO_2_. d) Box plot of phase transfer entropy (PTE) for both directions, where the box contains the interquartile range (IQR) and whiskers 1.5*IQR. On the left side, PTE between RESP_EEG_ and neural frequency bands. On the right side, PTE between ISF_EEG_ and RESP_EEG_. e) Probability density estimates of the average directional PTE (dPTE). Asterisks indicate statistical significance (colored: one-sample, black: two-sample, adjusted p<0.05). f) Topography shows median dPTE difference combined with permutation testing (p<0.05) whitening mask. Yellow circles around electrodes indicate significance with maximum statistic correction (p<0.05).

As described above, to investigate further the directional drive between the RESP_EEG_ and faster neural rhythms we quantified PTE and its directional form dPTE. Correlations were similar in both directions with respect to frequency band (Figure 3d). The baseline PTE magnitudes were clearly lower in comparison to the ISF-neural prediction. There was a robust dPTE between RESP_EEG_ and fast brain rhythms (Figure 3e) occurring at most every tested frequency band during wakefulness, except for the delta rhythm, which was unaltered by sleep. The slower RESP_EEG_ drove fast neural rhythms both during wakefulness (combined *dPTE*: 0.049 bits, *p_adj_*<0.002 for all, except for delta *padj*=0.189) and in sleep (mean *dPTE*: 0.085 bits, *p_adj_*<0.002, for delta band *p_adj_*<0.05). Topographical mapping (Figure 3f) separated occipital-parietal electrodes from the rest of the scalp, where there were only non-significant differences.

We also wanted to know if there was any interaction between the slow physiological ISF_EEG_ and RESP_EEG_. Results indicated increased phase-amplitude coupling during sleep with a slower ISF band (Figure 3a). Comparison against surrogate data revealed that this coupling was not widespread, being present at only about 10% of electrodes. Nonetheless, we still found a significant drive directed from ISF_1-EEG_ to RESP_EEG_ (Figure 3e). During sleep, the ISF_1-EEG_ drive of respiratory frequencies dropped significantly across most of the scalp, excepting some frontal electrodes (Figure 3f).

Taken together, these results confirmed that respiration phase couples with cortical rhythms, on a scale even larger than for the ISF. We further showed that respiration also drives neuronal amplitudes, whereas sleep state did not influence coupling strength, nor did it mediate changes in net directionality of this interaction, even though respiration frequency slowed down while showing increased spectral power.

## Discussion

In this electrophysiological study of healthy humans across awake and sleeping states, we have uncovered multiple novel interactions with directional coupling, namely infra-slow physiological oscillations, respiration, and fast cortical activity. We established that the phases of ISFs and respiration were mutually coupled and together exhibited directional phase-amplitude modulation of the amplitudes of fast neuronal activities over a wide range of frequencies during both wakefulness and sleep. These slow modulations of fast activities were independent of the power of the slow oscillations, and were obtained through phase-amplitude coupling. Moreover, we found that a transition in arousal state from wakefulness to sleep influenced this interaction, with a lowered net drive. Our findings thus show that the amplitudes of fluctuations of fast cortical oscillations are coordinated by the phase of slow and infra-slow physiological oscillations in a manner dependent on the arousal state.

### Cortical excitability is driven by the phase of ISFs

A comparison of wakefulness and sleep conditions showed that ISF_EEG_ power increased during sleep, along with a concurrent reduction in fast oscillation amplitudes (see Figure 1a). This result is well in line with previous findings that fluctuations in the BBB permeability to water increase during NREM sleep, insofar as scalp ISF_EEG_ potentials have been previously shown to reflect the potential difference across the BBB, namely between the blood compartment and the brain tissue interstitium (Voipio et al., 2003; Nita et al., 2004; Kiviniemi et al., 2017).

Previous work has shown that ISFs are phase-amplitude coupled with amplitudes of fast neuronal activities during detection-task performance (Monto et al., 2008) and sleep (Vanhatalo et al., 2004), albeit based on studies with limited electrode coverage and cohort sizes. We here used 256-channel fbEEG in a group of 21 subjects to corroborate and extend the earlier observations. We found that fast oscillation amplitudes were indeed phase-amplitude coupled with ISFs during wakefulness, and to an even great extent during sleep. Not only the magnitude but also the spatial extent wherein ISF_EEG_ phase coupled with neural amplitudes increased during sleep (see Figure 1 b-c). In conventional understanding, human sleep EEG is characterized by increased power of slow delta waves and the presence of sleep spindles occurring in the overlapping beta band (Erwin et al., 1984). In addition to these features of sleep, alpha rhythm is suppressed during the drowsiness preceding sleep (Erwin et al., 1984). Interestingly, the most conspicuous of the present ISF phase-amplitude coupling changes occurred in these same frequency bands, all of which are well-linked to sleep, thus strengthening the association between ISFs and sleep.

While these prior studies have demonstrated a phase-amplitude interaction of ISFs and respiration with fast neuronal activity, they relied upon correlative, undirected measures, which do not support drawing casual inferences. To resolve whether ISFs modulate the fast activities or if ISFs are rather a consequence of the emergent slow dynamics of fast activities, we used a novel directional phase-amplitude coupling analysis. In this approach, phase transfer entropy (PTE) quantifies in a history-controlled manner the mutual predictive powers of, respectively, ISF phase, and amplitudes of fast neural rhythms. Using PTE, we found that the ISF_EEG_ phase was a robust driver of the amplitude fluctuations of fast oscillations. Moreover, this directional phase-amplitude coupling was frequency dependent, such that the slowest frequencies (ISF_1-EEG_>ISF_2-EEG_>RESP_EEG_) were the strongest drivers.

One might reasonably expect the directional drive to be stronger during sleep than in wakefulness, given the precedents set by the circumstance of undirected phase-amplitude coupling and ISF power. However, the present analysis revealed that net dPTE drive was lower in sleep, to a similar degree in the fast→slow and slow→fast directions. These reductions were not linearly related to phase locking (see Figure S5), autocorrelation (see Figure S4), or bidirectionality of the interaction (see Figure 2a). In addition, we do not expect amplitude differences to be the cause since PTE is phase-based metric.

By exclusion, we suppose that the directed drive may thus arise through mechanisms that are partially independent from the pathways that achieve the instantaneous, correlative phase-amplitude coupling and that establish the overall signal power levels. This proposal seems to require invoking a new model in which a currently undefined driver could control the phase transitions both of ISF and neural oscillations during sleep. Such a model might necessarily require including additional factors such as brain water dynamics (Myllylä et al., 2018; Borchardt et al., 2021).

### Respiration phase operates on wide spatial extent during wakefulness and sleep

Strong respiratory brain pulsations have also been detected in human brain both using ultrafast MREG BOLD studies (Dreha-Kulaczewski et al., 2015, 2017; Kananen et al., 2020) and with intracranial needle electrode EEG (Zelano et al., 2016; Herrero et al., 2018) recordings. A recent study in awake mice revealed respiration-entrained oscillation patterns that were pacing electrical oscillations in the prefrontal cortex (Biskamp et al., 2017). Those findings inspired us to explore further whether respiration could in fact influence neurophysiological rhythms by affecting the ISF drive in human brain, either directly or indirectly.

Our results indicate that fast neuronal activity rhythms were not only phase-amplitude coupled, but were also driven by the respiration-related RESP_EEG_ phase, and not exclusively by the ISF_EEG_. Interestingly, unlike with ISF_EEG_, the phase-amplitude coupling and directional drive did not significantly differ between sleep and waking, and indeed the net drive tended to increase in the sleep state relative to waking. Furthermore, this net drive had a greater correlation with coupling values, which stands in contrast to ISF_EEG_ (see Figure S5).

The rate of CSF flow in mouse brain increases with the transition from waking to a sleep-like state with elevated slow-wave EEG characteristics (Ding et al., 2016). It may follow that the increased slow-wave power (EEG_SWS_ 0.2-2 Hz) that we detected over posterior regions during NREM sleep (see Figure 1a) could mark the zones of especially increased interstitial fluid volume and CSF movement. Previous studies have reported slow-wave EEG power increases to be most pronounced in frontal cortex (Finelli et al., 2000, 2001), however with different frequency band. A recent study ultrafast MREG BOLD study in humans showed that the areas of increased EEG slow-wave power during sleep also showed increases of all three physiological brain pulsations, especially the vasomotor and respiratory pulsations, and to a lesser extent also the cardiovascular pulsation increase (Helakari et al., 2022).

In light of this, we suppose that the physiological pulsations could have a causal relationship with diurnal neurophysiological changes. Previous work involving full night fMRI recordings with slow image sampling rate showed that < 0.1 Hz vasomotor waves increase in sleep (Chang et al., 2016; Liu et al., 2018). The several physiological pulsations affecting both blood and CSF dynamics could in theory be a unifying external factor to explain the reduced drive of neuronal activity by the electrical gradient across the BBB potential during sleep. This possible interaction requires further investigation. As an index of BBB permeability (Voipio et al., 2003; Nita et al., 2004) EEG ISFs may emerge as an object of study in a variety of brain injury and disease contexts. The generalization of this concept calls for further investigation of ISF dynamics with cortical rhythms in various clinical conditions.

### Limitations directing Future work

BBB permeability scans in conjunction with fbEEG recordings might eventually confirm the occurrence of glia limitans leakage in sleep. However, there is yet no technology for sufficiently fast recordings of BBB permeability changes across the sleep-wake cycle. Indeed, current experimental approaches are limited to invasive intracranial procedures that are apt to perturb the flow of interstitial fluid and CSF (Shoffstall et al., 2018; Plog et al., 2019). A non-invasive, multimodal neuroimaging investigation might eventually establish whether physiological CSF/blood pulsations are introducing hitherto undetected driving effects on sleep-related electrophysiological activity changes.

The awake and sleep scans were performed at different times of the day, such that circadian rhythms could have had an effect in the study. Our study design had limited recording time, and therefore describes the light sleep condition. For deeper sleep stages and REM-sleep, longer scan times would be necessary. After excluding various measurements, six of the 21 volunteers ended up providing unpaired measurements. We dealt with this potential confounding factor by using only unpaired testing.

## Materials and Methods

### Experimental Design

The study was approved by The Regional Ethics Committee of the Northern Ostrobothnia Hospital District. Written, informed consent was obtained from participants according to the Declaration of Helsinki. All the subjects were healthy non-smokers, with no continuous medication, neurological or cardio-respiratory diseases.

Thirty subjects were scanned twice in the study (Helakari et al., 2022) using multimodal imaging setup (Korhonen et al., 2014). One session with eyes fixated on a cross was obtained at 4-6 PM on the day after a full night of sleep, and another scan starting at 7-9 AM following a night of sleep deprivation. Each session consisted of two scans each lasting 10-15 minutes. Sleep deprivation was intended to enable the subjects to enter more quickly a deeper sleep state during the recordings (Kaufmann et al., 2006; Horovitz et al., 2009). Subjects were instructed not to drink any caffeinated beverages in the four hours before the awake resting state scan, and were requested to abstain for eight hours prior to the sleep deprivation scans. Consumption of alcohol was also prohibited in these intervals. Three subjects who, according to their sleep scores, did not fall asleep during the sleep deprived session, were excluded from the study (Table S1). Three subjects were also excluded due to suspicion of sleep apnea. Recordings were also disregarded in cases of insufficient data quality. Final group sizes were 21 awake (mean (SD) age 29.2 ± 6.8 years, 8 females) and 21 (28.4 ± 6.3 years, 11 females) sleep subjects.

FbEEG was recorded with the GES 400 (Electrical Geodesics) magnetic resonance imaging (MRI)-compatible system, with a 256-channel high density net (HydroCel Geodesic Sensor MR net), and DC-coupled amplifier (Net Amps 400). We used a sampling rate of 1 kHz (250 Hz for three sleep and five awake subjects, due to human error). The “Cz” electrode was used as a reference channel in the recordings. Signal quality and electrode impedances were inspected before the recordings. End-tidal carbon dioxide (ETCO_2_) was also measured in synchrony with the EEG and fMRI.

### Preprocessing

Gradient artifacts arising from switching of the MRI gradients were removed using template subtraction with Brain Vision Analyzer (v.2.1, Brain Products) (Allen et al., 2000). Template subtraction was also used to remove ballistocardiographic artefacts (Allen et al., 1998). The remaining signal processing steps and analyses were performed in Matlab (v.R2018b-2019b, MathWorks). All recordings were segmented to a matching duration of ten minutes. Linear trends were removed to attenuate stable baseline drift caused by conductivity changes at the electrode skin interface (Huigen et al., 2002), which is known to affect performance of various processing steps. We used independent component analysis (ICA) (FastICA (Hyvärinen, 1999)) along with principal component analysis dimension reduction (150 components) to transform datasets into independent components, in which artifactual ICs were identified and removed. Our main concern here was to remove ocular components, which can manifest as low (<1 Hz) frequency events. In the awake scans, subjects were instructed to fixate on a cross, which is known to reduce the frequency of saccades. To further minimize non-linearities and enhance the performance of ICA, we used a custom spike detection algorithm to remove artifactual spikes of very high amplitude. In place of the removed spikes, we interpolated the trends for the resultant gaps and used data from intact signal sequences to fill the gaps with real data imitating injections, in a process resembling inpainting (de Cheveigné and Arzounian, 2018). Bad channels were excluded from the ICA, followed by spherical interpolation. Recordings were referenced to linked mastoid electrodes, which were located near other electrodes, but which record less brain activity. Recordings were sleep-scored manually by experienced clinical neurophysiologists (JP, MK) following American Academy of Sleep Medicine guidelines (Table S1).

### Spectral analysis

Time-frequency (TF) spectral analysis (Figure 1a) was made with wavelet convolution using mirrored time-series to avoid edge effects. Wavelet convolution, implemented in the frequency domain, was made using complex Morlet wavelets (Eq.1). Conventional Fourier transform assumes signal stationarity, meaning that statistics of the signal, including mean, variance and frequency structure of the signal, do not change over time (Cohen, 2014). However, the stationarity assumption is violated by long EEG data, leading to decreased accuracy of frequency estimates. This, along with ability to generate time-resolved frequency representations with computational efficiency, led us to use wavelets. Fourier transform and Morlet wavelets both use sine waves as kernels, but in latter case the sine wave is not the full length of the recording and is tapered by a gaussian window (Cohen, 2014). The gaussian window is preferred to a rectangular window due to its smooth transition edges. The sharp edges in simple rectangular windowing give rise to artifactual ripple effects (Cohen, 2019).

Here, was used a logarithmic frequency range between 0.008 and 100 Hz, with 70 steps and constant wavelet cycle of 7, which is related to the width of the Gaussian by (Eq. 2). TF-power estimates were acquired from the squared magnitude of the convolution results. We used sleep scores to categorize the epoched TF-power estimates to episodes of wakefulness and sleep (N1-N3). These sleep scores served to discard awake epochs from the sleep samples, and vice versa, thus affording even more accurate spectral estimates. We then averaged over the time dimension and then power was formulated to relative amplitudes (Eq. 3) to reduce individual variability in power. For relative band power topographies, the power was summed over corresponding frequencies and divided by the total power over all recorded frequencies.

### Phase-amplitude coupling

We designed finite impulse response (FIR) Hamming windowed bandpass filters to filter the ISF into two separate bands (ISF_1-EEG_: 0.008-0.05 Hz & ISF_2-EEG_: 0.05-0.1 Hz). The ISF range of 0.008-0.1 Hz was chosen to obtain maximum coverage, without crosstalk with respiratory frequencies. Here we used a minimum filter kernel length defined by one cycle of the lowest frequency of interest. To avoid edge artifacts otherwise arising in the temporal filtering process, the signal was mirrored from both ends. These mirrored time-series *χ*(*n*),(*n* = 1,&,*N*) were zero-phase filtered *χ_ISF_*(*n*), to prevent phase distortions and offsets. FIR bandpass filters were designed similarly to five higher frequency EEG bands (delta: 1-4 Hz, theta: 4-8 Hz, alpha: 8-13 Hz, beta: 13-30 Hz, gamma: 30-40 Hz) using filter kernel length of 6 times the low cutoff frequency period to increase stopband attenuation, and zero-phase filtered to produce *χ_fast_*(*n*) signals.

Phase-amplitude coupling is a form of cross-frequency coupling where the phase of the slower rhythm is coupled to the amplitude of the faster oscillation. We used PLV (Lachaux et al., 1999) to assess the magnitude of coupling between the ISF phase and fast amplitudes, following methodology from (Vanhatalo et al., 2004; Palva et al., 2005; Monto et al., 2008). A Hilbert transform was applied, thus producing analytical signals *z_ISF_*(*n*) (Eq.4), from which the ISF phase time-series were extracted as *θ_ISF_* = *arg*(*z_ISF_*(*n*)). For faster frequency bands, the Hilbert amplitude envelopes were computed from the complex magnitude of the Hilbert transforms *a*(*n*) = |*z_fast_*(^*n*^)|. Amplitude envelopes were then filtered to infra-slow frequency bands using the same filters as used previously to produce *A_fast_*(*n*). Since phase-amplitude coupling operates on temporal resolution of the infra-slow phase, we down-sampled signals to 3 Hz to speed up computation. The Hilbert transform was applied to envelopes *A_fast_*(*n*) to extract the instantaneous phase *θ’_fast_* = *arg*(*z_A__fast_*(*n*)). Phase-amplitude coupling was quantified as 1:1 phase locking between *θ_ISF_* and *θ’_fast_*. PLV (Eq-5) was calculated for each electrode between both ISF bands and faster frequency bands. We present a diagram of the workflow in Supplementary Figure S1.

The respiratory frequency phase coupling with neural oscillations amplitudes was calculated similarly as with ISF. Individual respiratory frequencies were calculated from simultaneous end-tidal CO_2_ recordings using short-time Fourier transform with 50 second Hamming-tapered windowing and 50% overlap. The peak of the respiration frequency was then used as a center frequency for defining the complex Morlet wavelet (N=5). Wavelet convolution implemented in the frequency domain was then used to filter the recordings to respiratory frequencies, as follows: we compared RESP_EEG_ with neural bands, which were handled similarly as with the ISFs, with the distiniction that the envelopes were now filtered down to respiratory frequency. Furthermore, we compared RESP_EEG_ to ISFs phase-amplitude coupling using the same methods and filters as described above.

### Phase transfer entropy

We calculated directional predictive analyses using PTE (Lobier et al., 2013), which describes information flow between phases of slow physiological rhythms and faster neuronal rhythm amplitudes, i.e., determining whether an observation of the source helps to predict the phase transitions of the target signal. PTE is generally used in the same fashion as real-valued transfer entropy, but is instead applied to instantaneous phase signals.

We used the previously acquired instantaneous phases *θ_slow_* and *θ’_fast_* in the subsequent calculations, applying a 125 Hz sampling rate. Shannon’s entropy was used to calculate that reduction in uncertainty, which in this context defines the information content. State space transition of the data was used to produce the discrete probability distributions, from which Shannon entropies were calculated. Scott’s choice (Scott, 1979) was used to estimate the number of bins (Eq. 6). For PTE computation (Eq. 7), we calculated the probabilities for joint entropies and individual entropies in equations 8-10). Log base 2 was used to express in units of bits (Bossomaier et al., 2016; Timme and Lapish, 2018). We then formulated the PTE values into its directional form, dPTE (Eq.11), such that the sign of the metric indicates the direction of net information flow. PTE requires an *a priori* guess of the analysis lag, which we set to one cycle of the slow phase. However, PTE values are not highly sensitive to the analysis lag, such that connectivity can be accurately detected over a wide range of lags (Lobier et al., 2013).

### Statistical analysis

No statistical methods were used to pre-determine sample sizes. The null hypothesis power difference distribution (Figure 1a) was constructed with randomization testing using 10,000 permutations of shuffling of the group labels, where the null hypothesis held that vigilance states have the same effect on spectral power (two-tailed). The differential power map was then transformed to z-scores with respect to the permuted difference maps (Eq.12), for a metric of statistical significance. We assessed the multiple comparison problem using a maximum statistic correction, in which the null distribution was built from minimum and maximum values of the permuted maps, and the statistical significance threshold was extracted over the 2.5-97.5 percentile range.

Median PLVs were compared groupwise (Figures: 1b, left & 3a, left) using the Wilcoxon rank sum test with false discovery rate adjustment (Benjamini and Hochberg, 1995). We tested the null-hypothesis that vigilance state had no effect on phase-amplitude coupling (two-tailed). For average PLV difference topographies and dPTE topographies (Figures 1c, 2c, 3b, 3f) we permuted subject labels 10,000 times to build a null distribution, from which we acquired the first significance threshold criteria (again with thresholds between 2.5 and 97.5 percentiles). Maximum statistic correction was used to create even more strict significance criteria.

To assess the significance of the phase-amplitude coupling on the individual level (Figures 1b, right & 3a, right), we used time shifted surrogate data (Theiler et al., 1992; Lachaux et al., 1999). Null hypothesis distributions were built by splitting the phase of time-series x(t=1,…,T) from a random time-point k into x_1_=x(1,…,k) and x_2_=x(k,…,T), thus constructing the surrogate timeseries defined as xs=[x2,x1]. This approach preserves the temporal autocorrelation structure in the data. We shuffled the data 100 times to produce null-distribution of surrogate PLVs from which p-values were obtained (Eq.13). In all the analysis, the adjusted significance threshold was set as p<0.05 for rejection of the null hypothesis.

We tested whether the median dPTE differed from zero, which would indicate the presence of directionality in the phase-amplitude interaction. Here, we applied the one sample sign test with FDR-correction (Benjamini-Hochberg) to adjust p-values with 95% confidence level criteria (Figures 2b & 3e). The two sample Wilcoxon rank sum test with FDR correction was used to compare the medians between awake and sleep groups (Figures 2b & 3e). Our null hypothesis was that the medians of the two were equal (two-tailed), with the adjusted significance threshold at again p<0.05. Effect size estimates (η^2^) were calculated from z-values using the formula: 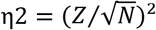, where N equals the total sample size (Fritz et al., 2012).

## Acknowledgments

This work was funded by Jane & Aatos Erkko Foundation grants 1&2 (V.Ki.), Academy of Finland Terva grants # 275342, 314497, 335720 (V.Ki), Aivosäätiö/BrainFoundation (J.K, V.Ki), Uniogs/MRC Oulu Doctoral Program-grant (H.H., J.K.), Pohjois-Suomen Terveydenhuollon tukisäätiö (H.H., V.Ko.), Instrumentarium Science Foundation (J.K), Emil Aaltonen Foundation (H.H, M.J), Finnish Medical Foundation (V.Ki., J.K., M.J.), VTR grants from Oulu University Hospital (V.Ko.), Finnish Neurological Foundation (V. Ki.), KEVO grants from Oulu University Hospital (V.Ki.), Orion Research Foundation (J.K.), Maire Taponen Foundation (J.K.). We acknowledge Inglewood Biomedical Editing for text revisions, Palva lab for providing a code for PTE calculation and CSC for computational resources. We thank Jussi Kantola for data management and Annastiina Kivipää for assistance during measurements and volunteer recruitment.

## References

Allen PJ, Josephs O, Turner R (2000) A method for removing imaging artifact from continuous EEG recorded during functional MRI. Neuroimage 12:230–239.

Allen PJ, Polizzi G, Krakow K, Fish DR, Lemieux L (1998) Identification of EEG events in the MR scanner: the problem of pulse artifact and a method for its subtraction. Neuroimage 8:229–239.

Benjamini Y, Hochberg Y (1995) Controlling the False Discovery Rate: A Practical and Powerful Approach to Multiple Testing. Journal of the Royal Statistical Society Series B (Methodological) 57:289–300 Available at: https://www.jstor.org/stable/2346101.

Besson JM, Woody CD, Aleonard P, Thompson HK, Albe-Fessard D, Marshall WH (1970) Correlations of brain d-c shifts with changes in cerebral blood flow. Am J Physiol 218:284–291.

Biskamp J, Bartos M, Sauer J-F (2017) Organization of prefrontal network activity by respiration-related oscillations. Sci Rep 7:45508 Available at: https://www.ncbi.nlm.nih.gov/pubmed/28349959.

Borchardt V, Korhonen V, Helakari H, Nedergaard M, Myllylä T, Kiviniemi V (2021) Inverse correlation of fluctuations of cerebral blood and water concentrations in humans. The European Physical Journal Plus 136:497.

Bossomaier T, Barnett L, Harré M, Lizier JT (2016) An Introduction to Transfer Entropy: Information Flow in Complex Systems. Springer International Publishing. Available at: https://www.springer.com/gp/book/9783319432212.

Cao Y, Roy S, Sachdev RNS, Heck DH (2012) Dynamic Correlation between Whisking and Breathing Rhythms in Mice. Journal of Neuroscience 32:1653–1659.

Ceguerra R v., Lizier JT, Zomaya AY (2011) Information storage and transfer in the synchronization process in locally-connected networks. In: 2011 IEEE Symposium on Artificial Life (ALIFE), pp 54–61. IEEE.

Chang C, Leopold DA, Schölvinck ML, Mandelkow H, Picchioni D, Liu X, Ye FQ, Turchi JN, Duyn JH (2016) Tracking brain arousal fluctuations with fMRI. Proceedings of the National Academy of Sciences 113:4518–4523.

Cohen MX (2014) Analyzing Neural Time Series Data. The MIT Press.

Cohen MX (2019) A better way to define and describe Morlet wavelets for time-frequency analysis. Neuroimage 199:81–86.

Cuddapah VA, Zhang SL, Sehgal A (2019) Regulation of the Blood-Brain Barrier by Circadian Rhythms and Sleep. Trends Neurosci 42:500–510.

de Cheveigné A, Arzounian D (2018) Robust detrending, rereferencing, outlier detection, and inpainting for multichannel data. Neuroimage 172:903–912.

Ding F, O’Donnell J, Xu Q, Kang N, Goldman N, Nedergaard M (2016) Changes in the composition of brain interstitial ions control the sleep-wake cycle. Science 352:550–555.

Dreha-Kulaczewski S, Joseph AA, Merboldt K-D, Ludwig H-C, Gärtner J, Frahm J (2015) Inspiration Is the Major Regulator of Human CSF Flow. Journal of Neuroscience 35:2485–2491.

Dreha-Kulaczewski S, Joseph AA, Merboldt K-D, Ludwig H-C, Gärtner J, Frahm J (2017) Identification of the Upward Movement of Human CSF In Vivo and its Relation to the Brain Venous System. Journal of Neuroscience 37:2395–2402.

Erwin CW, Somerville ER, Radtke RA (1984) A review of electroencephalographic features of normal sleep. J Clin Neurophysiol 1:253–274.

Finelli LA, Baumann H, Borbély AA, Achermann P (2000) Dual electroencephalogram markers of human sleep homeostasis: correlation between theta activity in waking and slow-wave activity in sleep. Neuroscience 101:523–529.

Finelli LA, Borbély AA, Achermann P (2001) Functional topography of the human nonREM sleep electroencephalogram. European Journal of Neuroscience 13:2282–2290.

Flexman JE, Demaree RG, Simpson DD (1974) Respiratory phase and visual signal detection. Percept Psychophys 16:337–339.

Fritz CO, Morris PE, Richler JJ (2012) Effect size estimates: Current use, calculations, and interpretation. J Exp Psychol Gen 141:2–18.

Fukunaga M, Horovitz SG, van Gelderen P, de Zwart JA, Jansma JM, Ikonomidou VN, Chu R, Deckers RHR, Leopold DA, Duyn JH (2006) Large-amplitude, spatially correlated fluctuations in BOLD fMRI signals during extended rest and early sleep stages. Magn Reson Imaging 24:979–992.

Fultz NE, Bonmassar G, Setsompop K, Stickgold RA, Rosen BR, Polimeni JR, Lewis LD (2019) Coupled electrophysiological, hemodynamic, and cerebrospinal fluid oscillations in human sleep. Science (1979) Available at: https://www.science.org/doi/abs/10.1126/science.aax5440.

Helakari H, Korhonen V, Holst SC, Piispala J, Kallio M, Väyrynen T, Huotari N, Raitamaa L, Tuunanen J, Kananen J, Järvelä M, Tuovinen T, Raatikainen V, Borchardt V, Kinnunen H, Nedergaard M, Kiviniemi V (2022) Human NREM Sleep Promotes Brain-Wide Vasomotor and Respiratory Pulsations. The Journal of Neuroscience 42:2503–2515.

Held D, Fencl V, Pappenheimer JR (1964) Electrical potential of cerebrospinal fluid. J Neurophysiol 27:942–959 Available at: https://journals.physiology.org/doi/abs/10.1152/jn.1964.27.5.942.

Herrero JL, Khuvis S, Yeagle E, Cerf M, Mehta AD (2018) Breathing above the brain stem: volitional control and attentional modulation in humans. J Neurophysiol 119:145–159 Available at: https://journals.physiology.org/doi/full/10.1152/jn.00551.2017.

Hiltunen T, Kantola J, Abou Elseoud A, Lepola P, Suominen K, Starck T, Nikkinen J, Remes J, Tervonen O, Palva S, Kiviniemi V, Palva JM (2014) Infra-Slow EEG Fluctuations Are Correlated with Resting-State Network Dynamics in fMRI. Journal of Neuroscience 34:356–362.

Horovitz SG, Braun AR, Carr WS, Picchioni D, Balkin TJ, Fukunaga M, Duyn JH (2009) Decoupling of the brain’s default mode network during deep sleep. Proceedings of the National Academy of Sciences - PNAS 106:11376–11381 Available at: https://agris.fao.org/agris-search/search.do?recordID=US201301653150.

Hughes SW, Lőrincz ML, Parri HR, Crunelli V (2011) Infra-slow (<0.1 Hz) oscillations in thalamic relay nuclei: basic mechanisms and significance to health and disease states. In, pp 145–162.

Huigen E, Peper A, Grimbergen C (2002) Investigation into the origin of the noise of surface electrodes. Med Biol Eng Comput 40:332–338.

Hyvärinen A (1999) Fast and robust fixed-point algorithms for independent component analysis. IEEE Trans Neural Netw 10 3:626–634.

Ito J, Roy S, Liu Y, Cao Y, Fletcher M, Lu L, Boughter JD, Grün S, Heck DH (2014) Whisker barrel cortex delta oscillations and gamma power in the awake mouse are linked to respiration. Nat Commun 5:3572.

Iwabe T, Ozaki I, Hashizume A (2014) The respiratory cycle modulates brain potentials, sympathetic activity, and subjective pain sensation induced by noxious stimulation. Neurosci Res 84:47–59.

Johannknecht M, Kayser C (2022) The influence of the respiratory cycle on reaction times in sensory-cognitive paradigms. Sci Rep 12:2586.

Kananen J, Helakari H, Korhonen V, Huotari N, Järvelä M, Raitamaa L, Raatikainen V, Rajna Z, Tuovinen T, Nedergaard M, Jacobs J, LeVan P, Ansakorpi H, Kiviniemi V (2020) Respiratory-related brain pulsations are increased in epilepsy—a two-centre functional MRI study. Brain Commun 2:fcaa076 Available at: https://search.proquest.com/docview/2444606699.

Kaufmann C, Wehrle R, Wetter TC, Holsboer F, Auer DP, Pollmächer T, Czisch M (2006) Brain activation and hypothalamic functional connectivity during human non-rapid eye movement sleep: an EEG/fMRI study. Brain 129:655–667.

Kiviniemi V, Korhonen V, Kortelainen J, Rytky S, Keinänen T, Tuovinen T, Isokangas M, Sonkajärvi E, Siniluoto T, Nikkinen J, Alahuhta S, Tervonen O, Turpeenniemi-Hujanen T, Myllylä T, Kuittinen O, Voipio J (2017) Real-time monitoring of human blood-brain barrier disruption. PLoS One 12:e0174072.

Kjaerby C, Andersen M, Hauglund N, Untiet V, Dall C, Sigurdsson B, Ding F, Feng J, Li Y, Weikop P, Hirase H, Nedergaard M (2022) Memory-enhancing properties of sleep depend on the oscillatory amplitude of norepinephrine. Nat Neurosci 25:1059–1070.

Kluger DS, Gross J (2021) Respiration modulates oscillatory neural network activity at rest. PLoS Biol 19:e3001457.

Korhonen V, Hiltunen T, Myllylä T, Wang X, Kantola J, Nikkinen J, Zang Y-F, LeVan P, Kiviniemi V (2014) Synchronous Multiscale Neuroimaging Environment for Critically Sampled Physiological Analysis of Brain Function: Hepta-Scan Concept. Brain Connect 4:677–689 Available at: https://doi.org/10.1089/brain.2014.0258.

Lachaux J-P, Rodriguez E, Martinerie J, Varela FJ (1999) Measuring phase synchrony in brain signals. Hum Brain Mapp 8:194–208 Available at: https://onlinelibrary.wiley.com/doi/abs/10.1002/%28SICI%291097-0193%281999%298%3A4%3C194%3A%3AAID-HBM4%3E3.0.CO%3B2-C.

Leopold DA, Murayama Y, Logothetis NK (2003) Very slow activity fluctuations in monkey visual cortex: implications for functional brain imaging. Cerebral Cortex 13:422–433.

Li S, Rymer WZ (2011) Voluntary Breathing Influences Corticospinal Excitability of Nonrespiratory Finger Muscles. J Neurophysiol 105:512–521.

Liu X, de Zwart JA, Schölvinck ML, Chang C, Ye FQ, Leopold DA, Duyn JH (2018) Subcortical evidence for a contribution of arousal to fMRI studies of brain activity. Nat Commun 9:395.

Lobier M, Siebenhühner F, Palva S, Palva JM (2013) Phase Transfer Entropy: A novel phase-based measure for directed connectivity in networks coupled by oscillatory interactions.

Marshall L, Mölle M, Fehm HL, Born J (1998) Scalp recorded direct current brain potentials during human sleep. Eur J Neurosci 10:1167–1178.

Marshall L, Molle M, Michaelsen S, Fehm HL, Born J (1996) Slow potential shifts at sleep--wake transitions and shifts between NREM and REM sleep. Sleep 19:145–151.

Monto S, Palva S, Voipio J, Palva JM (2008) Very Slow EEG Fluctuations Predict the Dynamics of Stimulus Detection and Oscillation Amplitudes in Humans. Journal of Neuroscience 28:8268–8272.

Myllylä T, Harju M, Korhonen V, Bykov A, Kiviniemi V, Meglinski I (2018) Assessment of the dynamics of human glymphatic system by near-infrared spectroscopy. J Biophotonics 11.

Niedermeyer E, Lopes da Silva, Fernando H, Donald L. Schomer, Vanhatalo S, Voipio J, Kaila K (2011) Electroencephalography: Basic principles, clinical applications, and related fields, 6th ed. Wolters Kluwer Health/Lippincott Williams & Wilkins.

Nita DA, Vanhatalo S, Lafortune F-D, Voipio J, Kaila K, Amzica F (2004) Nonneuronal origin of CO2-related DC EEG shifts: an in vivo study in the cat. J Neurophysiol 92:1011–1022.

Osorio-Forero A, Cardis R, Vantomme G, Guillaume-Gentil A, Katsioudi G, Devenoges C, Fernandez LMJ, Lüthi A (2021) Noradrenergic circuit control of non-REM sleep substates. Current Biology 31:5009–5023.e7.

Palva JM, Palva S (2012) Infra-slow fluctuations in electrophysiological recordings, blood-oxygenation-level-dependent signals, and psychophysical time series. Neuroimage 62:2201–2211.

Palva JM, Palva S, Kaila K (2005) Phase synchrony among neuronal oscillations in the human cortex. J Neurosci 25:3962–3972.

Palva JM, Zhigalov A, Hirvonen J, Korhonen O, Linkenkaer-Hansen K, Palva S (2013) Neuronal long-range temporal correlations and avalanche dynamics are correlated with behavioral scaling laws. Proceedings of the National Academy of Sciences 110:3585–3590.

Plog BA, Lou N, Pierre CA, Cove A, Kenney HM, Hitomi E, Kang H, Iliff JJ, Zeppenfeld DM, Nedergaard M, Vates GE (2019) When the air hits your brain: decreased arterial pulsatility after craniectomy leading to impaired glymphatic flow. J Neurosurg:1–14.

Scott DW (1979) On optimal and data-based histograms. Biometrika 66:605–610 Available at: https://doi.org/10.1093/biomet/66.3.605.

Shoffstall AJ, Paiz JE, Miller DM, Rial GM, Willis MT, Menendez DM, Hostler SR, Capadona JR (2018) Potential for thermal damage to the blood–brain barrier during craniotomy: implications for intracortical recording microelectrodes. J Neural Eng 15:034001.

Tétrault S, Chever O, Sik A, Amzica F (2008) Opening of the blood-brain barrier during isoflurane anaesthesia. European Journal of Neuroscience 28:1330–1341.

Theiler J, Eubank S, Longtin A, Galdrikian B, Doyne Farmer J (1992) Testing for nonlinearity in time series: the method of surrogate data. Physica D 58:77–94 Available at: https://www.sciencedirect.com/science/article/pii/016727899290102S.

Timme NM, Lapish C (2018) A Tutorial for Information Theory in Neuroscience. eNeuro 5.

Tschirgi RD, Taylor JL (1958) Slowly Changing Bioelectric Potentials Associated With the Blood Brain Barrier. American Journal of Physiology-Legacy Content 195:7–22.

Vanhatalo S, Palva JM, Holmes MD, Miller JW, Voipio J, Kaila K (2004) Infraslow oscillations modulate excitability and interictal epileptic activity in the human cortex during sleep. Proc Natl Acad Sci U S A 101:5053–5057 Available at: http://www.pnas.org/content/101/14/5053.abstract.

Voipio J, Tallgren P, Heinonen E, Vanhatalo S, Kaila K (2003) Millivolt-Scale DC Shifts in the Human Scalp EEG: Evidence for a Nonneuronal Generator. J Neurophysiol 89:2208–2214 Available at: https://doi-org.pc124152.oulu.fi:9443/10.1152/jn.00915.2002.

Zelano C, Jiang H, Zhou G, Arora N, Schuele S, Rosenow J, Gottfried JA (2016) Nasal Respiration Entrains Human Limbic Oscillations and Modulates Cognitive Function. Journal of Neuroscience 36:12448–12467.

