## Supplemental Information for "Sleep specific changes in infra-slow and respiratory frequency drivers of cortical EEG rhythms"

Supplementary information includes:

Supplementary information text
Tables S1 to S2
Figures S1 to S6
SI References

### Supplementary Information Text

#### Expanded methods

**Sleep classification.** Sleep scores (Table S1) were made by experienced clinical neurophysiologists who were trained for sleep scoring. The scorings were made in 30 second epochs following American Academy of Sleep Medicine guidelines for clinical sleep studies. The recordings used in the scoring were gradient and BCG-artifact corrected. EEG epochs were scored as awake, N1 (light sleep), N2 (intermediate sleep with K-complexes and/or sleep spindles), and N3 (slow wave sleep). Table S1 shows the scoring with '1-3' indicating the non-REM sleep depth, 'w' for wakefulness, and '-' for noise contaminated epochs, from which sleep scoring were not possible to assess.

**Phase-amplitude coupling workflow**. Paired recording was used to visualize the workflow of the calculations (Figure S1) on the left and right paired awake and sleep recordings. A 100 s segment is shown from electrode E160 in a representative example. The fbEEG trace $x(n)$ is filtered into ISF (<0.1 Hz), delta (1-4 Hz), theta (4-8 Hz), alpha (8-13 Hz), beta (13-30 Hz) and gamma (30-40 Hz) frequency bands. In the figure, only the slower ISF and related products are shown. Fast amplitude $x_{fast}(n)$ was then Hilbert transformed, from which the analytical envelopes were obtained. Envelopes were then filtered to infra-slow frequencies using same ISF filters as before to produce $A_{fast}(n)$. The Hilbert transform was applied again to acquire instantaneous phase-evolutions for the $A_{fast}(n)$ and $x_{ISF}(n)$. Using $\theta_{ISF}(n)$ and $\theta'_{fast}(n)$ phases we calculated PLV (Eq. 5) (Lachaux et al., 1999) over time, giving one phase locking value for the whole recording.

<u>Collection of formulas</u>

(Eq.1) $\omega = e^{i2\pi ft} e^{\frac{-t^2}{2\sigma^2}}$

(Eq.2) $\sigma = \frac{n}{2\pi f}$

(Eq.3) $RA_i = ({}^{A_i}/_{\sum_i A_i}) * 100\%$

(Eq.4) $z(n) = x(n) + iy(n) = a(n)e^{i\theta(n)}$

where n denotes to discrete time variable, $a(n)$ and $\theta(n)$ are the instantaneous amplitude and phase.

(Eq.5) $PLV_{slow,fast} = \frac{1}{N}\left|\sum_{n=1}^{N} e^{i(\theta_{slow}(n)-\theta'_{fast}(n))}\right|$

where N is the number of samples

(Eq.6) $k = \frac{2\pi}{h}, where\ h = \frac{3.5\sigma}{N^{\frac{1}{3}}}$

(Eq.7) $pTE_{x\to y} = H\left(\theta_y(n), \theta_y(n')\right) + H\left(\theta_y(n'), \theta_x(n')\right) - H\left(\theta_y(n')\right) - H(\theta_y(n), \theta_y(n'), \theta_x(n'))$

(Eq.8) $H(\theta_x(n)) = -\sum p(\theta_x(n)) \log_2 p(\theta_x(n))$

(Eq.9) $H\left(\theta_x(n'), \theta_y(n)\right) = -\sum p\left(\theta_x(n'), \theta_y(n)\right) \log_2 p\left(\theta_x(n'), \theta_y(n)\right)$

(Eq.10) $H\left(\theta_y(n), \theta_y(n'), \theta_x(n')\right) = -\sum p\left(\theta_y(n), \theta_y(n'), \theta_x(n')\right) \log_2 p\left(\theta_y(n), \theta_y(n'), \theta_x(n')\right)$

(Eq.11) $dpTE_{x\to y} = pTE_{x\to y} - pTE_{y\to x}$

(Eq.12) $z = \frac{x-\mu}{\sigma}$

(Eq.13) $p = N(PLV_{surrogate} > PLV)/N_{permutations}$

(Eq.14) $H\left(\theta_y(n), \theta_y(n')\right) = -\sum p\left(\theta_y(n), \theta_y(n')\right) \log_2 p\left(\theta_y(n), \theta_y(n')\right)$

**Supplementary results**

**$ISF_{2\text{-}EEG}$ phase-amplitude coupling and directional drive**. We obtained phase-amplitude coupling PLV and dPTE between $ISF_{2\text{-}EEG}$ and neural cortical rhythms in similar manner as with the $ISF_{1\text{-}EEG}$ presented in the main text methods.

We found similar coupling patterns with $ISF_{2\text{-}EEG}$ as with the slower $ISF_{1\text{-}EEG}$ band, where the coupling was stronger throughout all the frequencies during sleep (Figure S2a left). Coupling magnitudes were generally lower than with $ISF_{1\text{-}EEG}$. Significant differences between the two states were seen only with $ISF_{2\text{-}EEG}$-beta coupling. Surrogate testing confirmed that not only the coupling magnitudes, but also number of significantly coupled electrodes, were lower in this ISF range (Figure S2a right). Interquartile ranges of significantly phase-amplitude coupled electrodes were 5 to 20 percent, depending on the frequency band. Especially the sleep state showed a decrease in the extent of coupling in comparison to $ISF_1$. Significant differences in PLV between the two states were focused on delta and beta coupling in frontal areas (Figure S2b). Average phase difference probabilities showed only scant emphasis on small phase differences (Figure S2c). The phase differences between $ISF_2$ and fast oscillations were more dominant in range of 0 to π occurring at frontal electrodes during sleep.

Baselines of PTE values in both directions were lower that with $ISF_1$ and there seemed to be a stronger prediction from fast oscillations amplitudes towards slow ISF (Figure S3a). Reversed net dPTE was found to occur with $ISF_{2\text{-}EEG}$, directed from fast activity to slow ISF phase (Figure S3b). This was significant during wakefulness with delta, theta, and gamma frequency bands. However, the two-sample statistical testing showed no significant differences between groups. Spatial mapping of the dPTE differences (Figure S3c) between the arousal states confirmed the previous results, with only individual electrodes showing significant differences.

**Autocorrelation and PTE**. Increased spectral power of the low frequency bands during sleep could indicate the direction of increased autocorrelations. Phase transfer entropy (Lobier et al., 2013), even though it is connectivity measure, is ultimately affected by correlations. An increase in correlations could lead to decreased information transfer. We used autocorrelation in terms of transfer entropy (Eq. 14) to test the autocorrelation with the same analysis lag used in the PTE calculations (Figure S4). If the autocorrelations between the wakefulness and sleep differed significantly, it was not meaningful to compare the PTEs between conditions, since the differences could have arisen from differences in autocorrelations. An average was taken over all channels, and the Wilcoxon rank sum combined with FDR correction was used to assess statistical significance, with the null hypothesis being that the medians of the two distributions were equal. No significant differences were seen between the groups, suggesting that the comparison of dPTE values on a group level is valid, and that correlation differences do not explain the differences observed with PTE.

**Relation between PLV and PTE**. We wanted to test for a correlation between phase-amplitude coupling PLV and directional phase-amplitude coupling measured by dPTE magnitude. We supposed that increased coupling would be accompanied by increased information transfer, as is usually seen with Kuramoto models (Ceguerra et al., 2011). We took a median PLV and dPTE magnitude over subjects and combined all five cortical bands in one. Next, we performed linear regression analysis using ordinary least squares fitting (Figure S5). The coefficients of determination ($R^2$) between the model and observations were low, indicating that a linear model was not sufficient to explain variation in the observations. Residuals and residual autocorrelations showed that the model was not biased and contained no autocorrelation of residuals.

**Average phase differences between ISF phase and neural amplitudes**. Average phase-amplitude coupling phase differences (Figure S6) between ISF phase and all tested frequency bands, revealed that phase difference patterns were similar between neural frequencies. A transition from wakefulness to sleep was associated with a change whereby phase differences concentrated around pi/2, especially with $ISF_{1\text{-}EEG}$.

**Table for statistical tests**. We compiled all the statistical test results obtained in this study (Table S2), including phase-amplitude coupling PLV and dPTE, along with corresponding adjusted p-values and effect sizes for statistical tests described in the main manuscript.

**Table S1.** Sleep scores for individual EEG recordings along with final group sizes. Sleep was scored in 30 second epochs, where '1-3' indicates non-REM sleep depth, 'w' is wakefulness and '-' represents artifactual epochs. Each column corresponds to one epoch.

Sleep classifications

| Awake / epoch | 1 | 2 | 3 | 4 | 5 | 6 | 7 | 8 | 9 | 10 | 11 | 12 | 13 | 14 | 15 | 16 | 17 | 18 | 19 | 20 |
|---|---|---|---|---|---|---|---|---|---|---|---|---|---|---|---|---|---|---|---|---|
| Subject 1 | w | w | w | w | w | w | w | w | w | w | w | w | w | w | w | w | w | w | w | w |
| Subject 2 | w | w | w | w | w | w | w | w | w | w | w | w | w | w | w | w | w | w | w | w |
| Subject 3 | w | w | w | w | w | w | w | w | w | w | w | w | w | w | w | w | w | - | - | - |
| Subject 4 | w | w | w | w | w | w | w | w | w | w | w | w | w | w | w | w | w | w | w | w |
| Subject 5 | w | w | w | w | w | w | w | w | w | w | w | w | w | w | w | w | w | w | w | w |
| Subject 6 | w | w | w | w | w | w | w | w | w | w | w | w | w | w | w | w | w | w | w | w |
| Subject 7 | w | w | w | w | w | w | w | w | w | w | w | w | w | w | w | w | w | w | w | w |
| Subject 8 | w | w | w | w | w | w | w | w | w | w | w | w | w | w | w | w | w | w | w | w |
| Subject 9 | w | w | w | w | w | w | 1 | w | w | w | w | w | w | w | 1 | w | w | w | w | w |
| Subject 10 | w | w | w | w | w | w | w | w | w | w | w | w | w | w | w | w | w | w | w | w |
| Subject 11 | w | w | w | w | w | w | w | w | w | w | w | w | w | w | w | w | w | w | w | w |
| Subject 12 | w | w | w | w | w | w | w | w | w | w | w | w | w | w | w | w | w | w | w | w |
| Subject 13 | w | w | w | w | w | w | w | w | w | w | w | w | w | w | w | w | w | w | w | w |
| Subject 14 | w | w | w | w | w | w | w | w | w | w | w | w | w | w | w | w | w | w | w | w |
| Subject 15 | w | w | w | w | w | w | w | w | w | w | w | w | w | w | w | w | w | w | w | w |
| Subject 16 | w | w | w | w | w | w | w | w | w | w | w | w | w | w | w | w | w | w | w | w |
| Subject 17 | w | w | w | w | w | w | w | w | w | w | w | w | w | w | w | w | w | w | w | w |
| Subject 18 | w | w | w | w | w | w | w | w | w | w | w | w | w | w | w | w | w | w | w | w |
| Subject 19 | w | w | w | w | w | w | w | w | w | w | w | w | w | w | w | w | w | w | w | w |
| Subject 20 | w | w | w | w | w | w | w | w | w | w | w | w | w | w | w | w | w | w | w | w |
| Subject 21 | w | w | w | w | w | w | w | w | w | w | w | w | w | w | w | w | w | w | w | - |

| Sleep / epoch | 1 | 2 | 3 | 4 | 5 | 6 | 7 | 8 | 9 | 10 | 11 | 12 | 13 | 14 | 15 | 16 | 17 | 18 | 19 | 20 |
|---|---|---|---|---|---|---|---|---|---|---|---|---|---|---|---|---|---|---|---|---|
| Subject 1 | w | w | w | w | w | w | w | w | w | w/1 | w | 1 | 1 | 1 | 2 | 1 | 1 | 2 | 2 | 2 |
| Subject 2 | 1 | 1 | 1 | 1 | 2 | 2 | 1 | 1 | 2 | 2 | 2 | 2 | 2 | 2 | 1 | 1 | 1 | 1 | 1 | 2 |
| Subject 3 | 2 | 2 | w | w | 1 | 1 | 1 | 2 | 2 | 2 | 2 | 2 | 2 | 2 | 2 | 2 | 2 | 2 | 2 | 2 |
| Subject 4 | w | 1 | 1 | 1 | 1 | 1/2 | 1 | 1 | 1 | 1 | 1 | 1 | 1 | 1 | 1 | 1 | 2 | 2 | 2 | 1 |
| Subject 5 | w | w | w | 1 | w | w | 1 | w | 1 | 1 | 1 | w | w | w | w | w | w | 1 | w | w |
| Subject 6 | w | - | 1 | 1 | 1 | 1 | 2 | 2 | 2 | 1 | w | 2 | 2 | 2 | 2 | 2 | 2 | 2 | 2 | 2 |
| Subject 7 | w | w | w | 1 | 1 | 1 | 1 | 2 | 2 | 2 | 2 | 2 | 2 | 2 | 1 | 1 | 2 | 2 | 2 | 2 |
| Subject 8 | w | w | w | w | 1 | w | 1 | 1 | w | 1 | 1 | 1 | 1 | 1 | 1 | w | 1 | 2 | 1 | w |
| Subject 9 | 1 | 2 | 2 | 2 | 1 | w | 1/2 | w | 1 | w | w | 1 | 1 | 1 | 1 | w | 1 | 1 | 1 | 1 |
| Subject 10 | 2 | 2 | 2 | 2 | w | w | w | 1 | 2 | 2 | 2 | 2 | 2 | 2 | 1 | 2 | 1 | 2 | 2 | 3 |
| Subject 11 | 1 | 1 | 1 | 1 | 1 | - | 1 | 2 | 2 | - | 2 | 2 | 2 | 2 | 2 | 2 | 2 | 2 | 2 | - |
| Subject 12 | 1 | 2 | - | 1 | 2 | - | 2 | 2 | 2 | 1 | 2 | 2 | 2 | 2 | 2 | 1 | 1 | 2 | 1 | 2 |
| Subject 13 | w | w | 1 | 1 | 1 | 1 | w | 1 | 2 | 2 | w | w | w | 1 | 1 | w | w | w | w | w |
| Subject 14 | 1 | w | w | w | w | w | 2 | 2 | 2 | - | 1 | 2 | 2 | 2 | 2 | 2 | 1 | 2 | 2 | 2 |
| Subject 15 | w | w | 1 | 1 | 1 | 2 | 1 | 1 | 2 | 2 | 2 | 1 | 1 | 1 | w | 1 | 2 | 2 | 2 | 2 |
| Subject 16 | 1 | 1 | 1 | 1 | 1 | 1 | 1 | 2 | 2 | 1 | 1 | 2 | 1 | 1 | w | 1 | 1 | 2 | 1 | 1 |
| Subject 17 | 1 | 1 | 1 | w | w | w | - | 1 | 2 | 2 | w | 2 | 1 | 2 | 2 | 2 | 2 | w | 1 | 1 |
| Subject 18 | w | 1 | 1 | 1 | 1 | 1 | 1 | 1 | 1 | 1 | 1 | 1 | 1 | 1 | 1 | 1 | 1 | 1 | 1 | 1 |
| Subject 19 | w | 1 | 1 | 1 | 1 | w | w | 1 | 1 | 2 | 2 | 1 | 2 | 2 | 1 | 1 | 1 | 2 | 1 | 2 |
| Subject 20 | w | 1 | 1 | 1 | 2 | 2 | 1 | 1 | 1 | 2 | 2 | 2 | 1 | 1 | w | 1 | 1 | 2 | 2 | 1 |
| Subject 21 | w | w | w | w | w | 1 | 1 | 1 | 1 | 1 | 1 | 1 | 2 | 2 | 1 | 2 | 2 | 1 | 2 | 2 |

**Table S2.** Table of statistical tests. Median phase-locking values (PLV) and directional phase transfer entropy (dPTE) are shown, including adjusted p-values and effect sizes for 1- and 2-samples tests. Red color indicates awake state and blue colors sleep state. Significant p-values are shown bolded with 95% confidence criteria.

| | | Alpha | Beta | Theta | Delta | Gamma |
|---|---|---|---|---|---|---|
| **PLV** | **$ISF_1$** | 0.161 | 0.185 | 0.191 | 0.186 | 0.164 |
| | **$ISF_2$** | 0.155 | 0.150 | 0.165 | 0.167 | 0.152 |
| | **$ISF_1$** | 0.242 | 0.208 | 0.208 | 0.215 | 0.182 |
| | **$ISF_2$** | 0.170 | 0.171 | 0.176 | 0.188 | 0.160 |
| **2-sample $p_{adj}$** | **$ISF_1$** | **0.002** | **0.037** | 0.111 | **0.031** | 0.185 |
| | **$ISF_2$** | 0.164 | **0.037** | 0.164 | 0.069 | 0.563 |
| **Eff. Size $\eta^2$** | **$ISF_1$** | 0.339 | 0.142 | 0.080 | 0.179 | 0.046 |
| | **$ISF_2$** | 0.054 | 0.142 | 0.058 | 0.106 | 0.008 |
| **dPTE** | **$ISF_1$** | 0.110 | 0.090 | 0.093 | 0.099 | 0.103 |
| | **$ISF_2$** | -0.039 | -0.016 | -0.038 | -0.035 | -0.016 |
| | **$ISF_1$** | 0.062 | 0.053 | 0.052 | 0.038 | 0.054 |
| | **$ISF_2$** | -0.026 | -0.015 | -0.005 | -0.003 | -0.012 |
| **2-sample $p_{adj}$** | **$ISF_1$** | **0.003** | **0.003** | **0.018** | **0.003** | **0.001** |
| | **$ISF_2$** | 0.217 | 0.960 | 0.051 | 0.218 | 0.349 |
| **Eff. Size $\eta^2$** | **$ISF_1$** | 0.271 | 0.259 | 0.163 | 0.243 | 0.376 |
| | **$ISF_2$** | 0.049 | 6.03E-05 | 0.111 | 0.044 | 0.024 |
| **1-sample $p_{adj}$** | **$ISF_1$** | **4.77E-06** | **5.25E-05** | **5.25E-05** | **4.43E-04** | **4.77E-06** |
| | **$ISF_2$** | **0.002** | 0.078 | **0.010** | **0.030** | **0.030** |
| | **$ISF_1$** | **0.001** | **0.004** | **9.54E-06** | **1.05E-04** | **0.014** |
| | **$ISF_2$** | 0.426 | 0.426 | 0.426 | 0.664 | 0.426 |

| | | Alpha | Beta | Theta | Delta | Gamma |
|---|---|---|---|---|---|---|
| **PLV** | **Resp** | 0.169 | 0.202 | 0.214 | 0.197 | 0.143 |
| | **Resp** | 0.173 | 0.163 | 0.161 | 0.173 | 0.166 |
| **2-sample $p_{adj}$** | **Resp** | 0.546 | 0.546 | 0.291 | 0.291 | 0.511 |
| **Eff. Size $\eta^2$** | **Resp** | 0.009 | 0.009 | 0.070 | 0.056 | 0.020 |
| **dPTE** | **Resp** | 0.060 | 0.041 | 0.047 | 0.051 | 0.048 |
| | **Resp** | 0.096 | 0.080 | 0.093 | 0.073 | 0.084 |
| **2-sample $p_{adj}$** | **Resp** | 0.597 | 0.597 | 0.597 | 0.597 | 0.597 |
| **Eff. Size $\eta^2$** | **Resp** | 0.007 | 0.007 | 0.047 | 0.008 | 0.023 |
| **1-sample $p_{adj}$** | **Resp** | **4.43E-04** | **4.43E-04** | **0.002** | 0.189 | **0.002** |
| | **Resp** | **0.002** | **4.43E-04** | **2.10E-04** | **0.030** | **4.43E-04** |

| | | $ISF_1$ | $ISF_2$ |
|---|---|---|---|
| **PLV** | **Resp** | 0.167 | 0.160 |
| | **Resp** | 0.197 | 0.182 |
| **2-sample $p_{adj}$** | **Resp** | **0.008** | 0.349 |
| **Eff. Size $\eta^2$** | **Resp** | 0.251 | 0.039 |
| **dPTE** | **Resp** | 0.089 | 0.009 |
| | **Resp** | 0.046 | 0.017 |
| **2-sample $p_{adj}$** | **Resp** | **2.70E-05** | 0.125 |
| **Eff. Size $\eta^2$** | **Resp** | 0.451 | 0.056 |
| **1-sample $p_{adj}$** | **Resp** | **3.81E-06** | 0.078 |
| | **Resp** | **4.20E-05** | 0.078 |

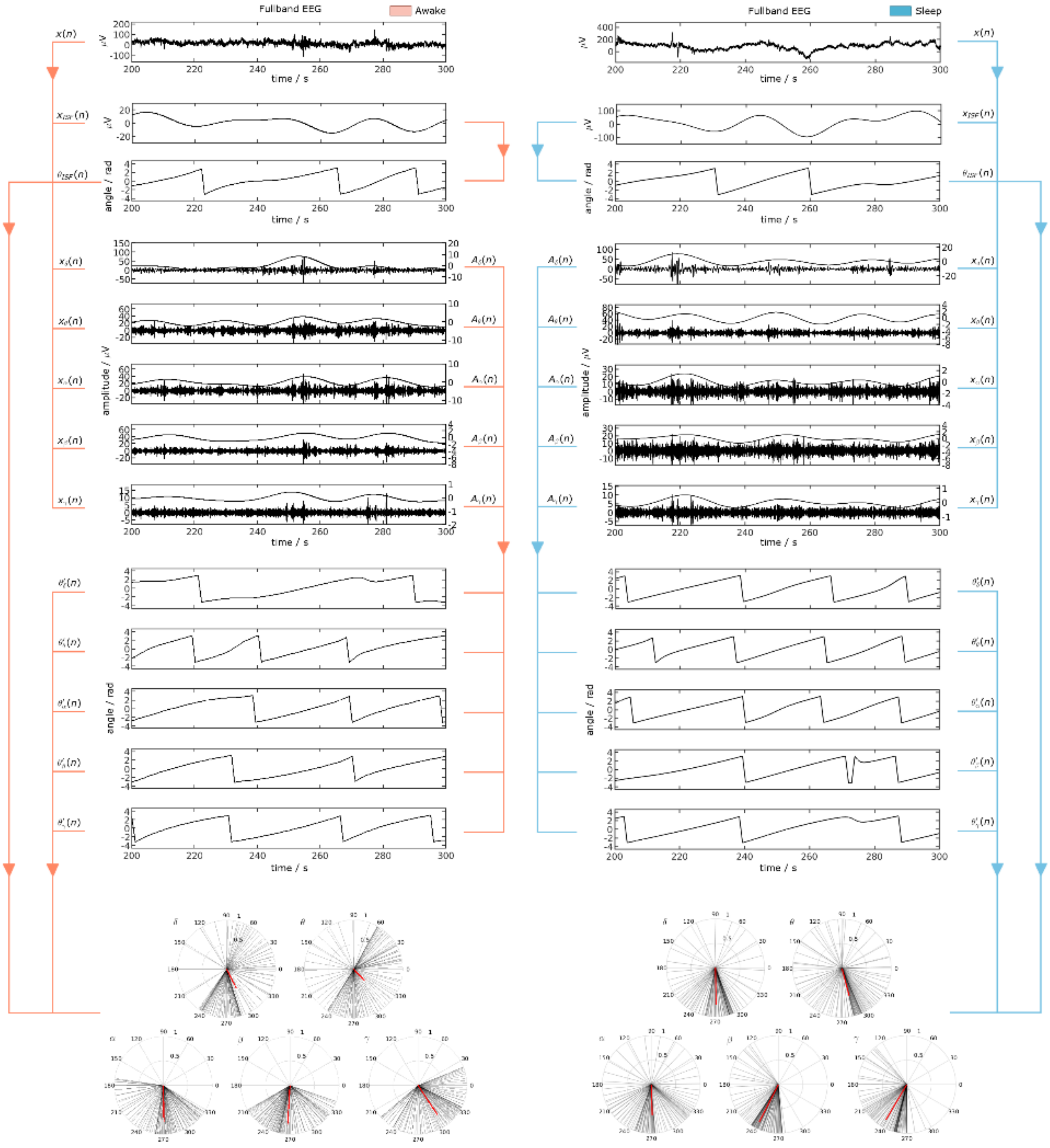


**Figure S1.** Workflow of the phase-amplitude coupling analysis with paired recording from one representative electrode. Arrows represent the calculation ordering. Unit circles are used to visualize the coupling pattern, where the length of the red line indicates phase-locking value (PLV).

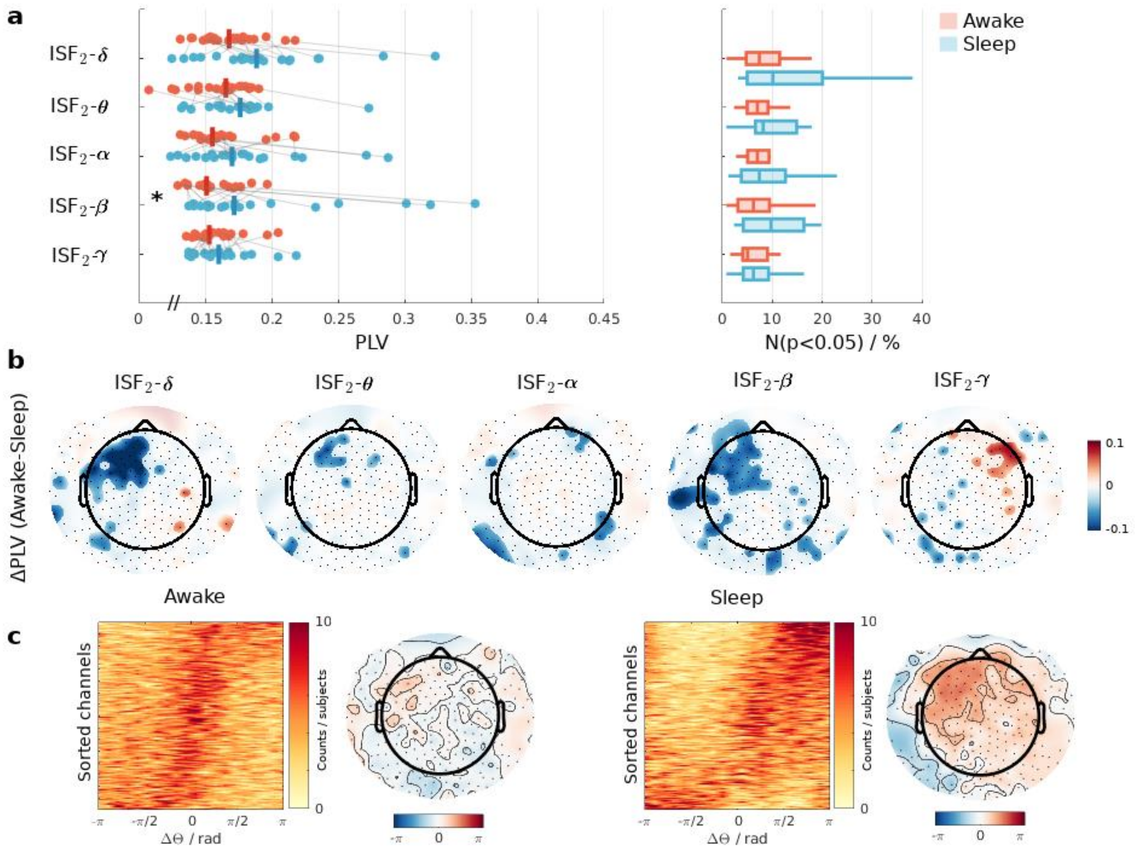


**Figure S2.** ISF2 phase-amplitude coupling with neural oscillations. Panel a) left: Median phase-locking value (PLV) taken over all electrodes. Asterisks indicate statistically significant differences in the coupling strength between awake and sleep. Gray lines connect paired test subjects. Right: The number of significantly coupled electrodes for each subject, scaled to percentiles. Panel b) The difference in average PLV combined with an overlayed significance mask ($p<0.05$). Panel c) Probability estimates of the average phase difference between ISF phase and faster rhythm phases. Channels on the y-axis are sorted into ascending order according to the median phase difference. Topographical plot shows the median phase difference taken over the five bands and for all subjects.

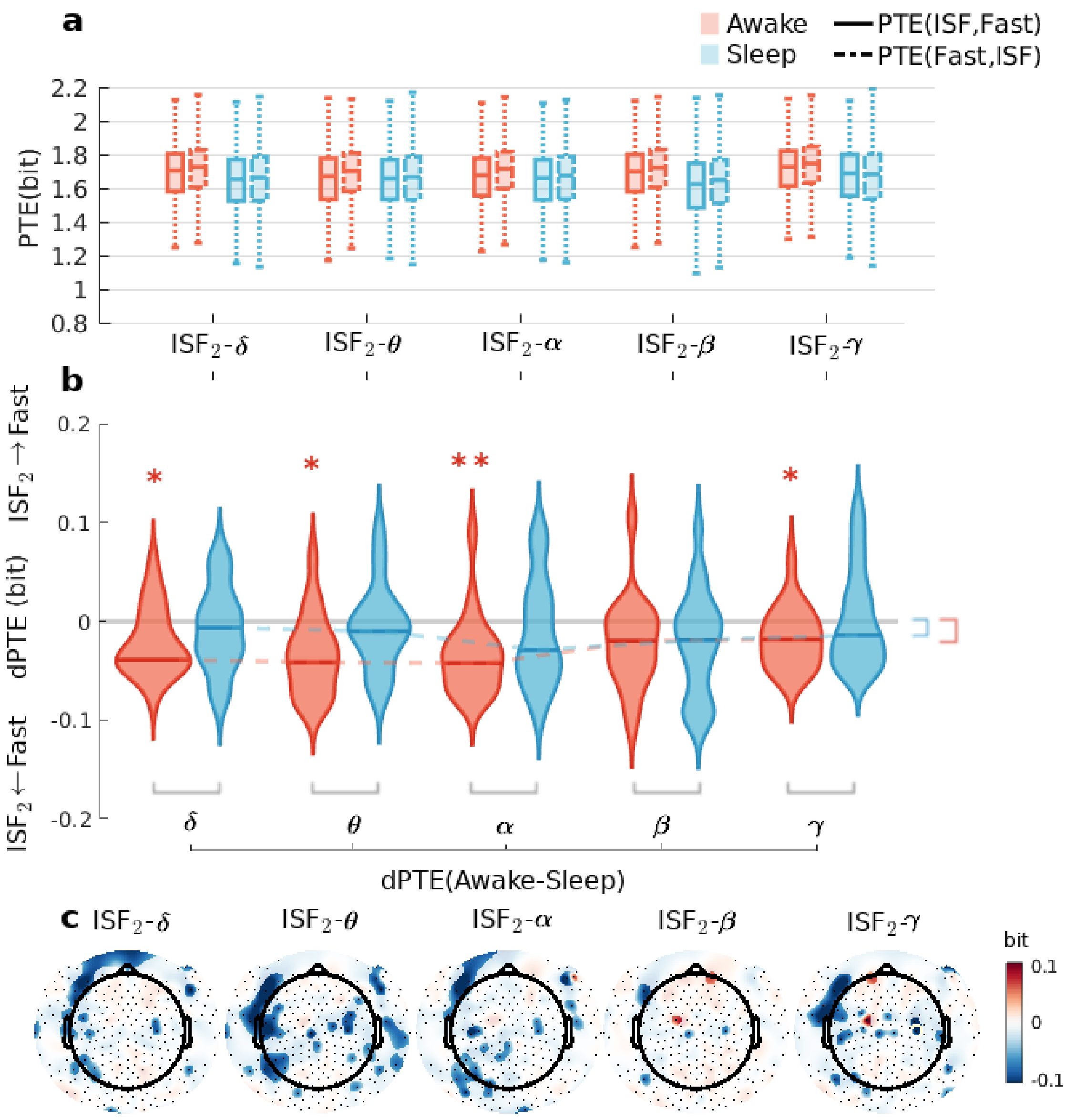


**Figure S3.** $ISF_2$ and neural oscillation directional drive. a) Box-plot of phase transfer entropy (PTE) magnitudes calculated in both directions. b) Probability density estimate of the average directional PTE (dPTE). Greek letters indicate the neural band in question. Asterisks indicate statistical significance (adjusted $p<0.05$). Colored asterisks are for one sample tests, indicating whether there is significant non-zero information transfer. Black asterisks are for two-sample tests i.e., groupwise comparison. c) Topographical presentation of average dPTE difference (awake - sleep) combined with overlayed significance mask ($p<0.05$). Yellow circles indicate the electrodes that were significant after maximum statistic correction ($p<0.05$).

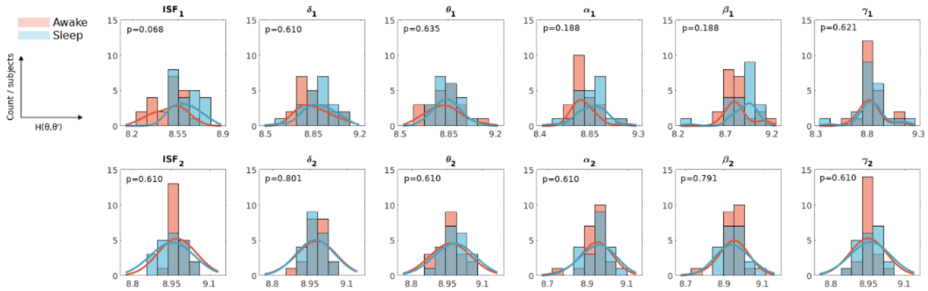

**Figure S4.** Differences in phase transfer entropy were not explained by differences in autocorrelations. The top row corresponds to slower $ISF_{1\text{-}EEG}$ (0.008-0.05 Hz) band and related products, and the lower row depicts the faster $ISF_{2\text{-}EEG}$ (0.05-0.1 Hz) band. The x-axis indicates autocorrelation score, and the y-axis depicts the histogram count. Greek letters represent slow filtered amplitude envelopes used in coupling analyses. P-values are false discovery rate (FDR) adjusted p-values.

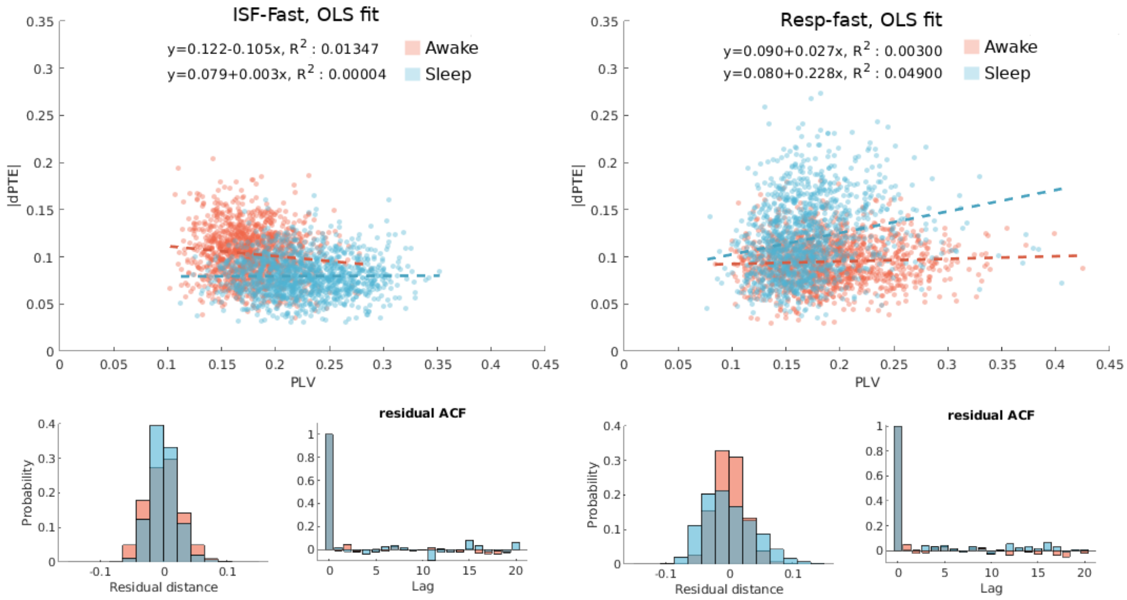


**Figure S5.** Magnitude of information transfer as a function of phase-locking value for $ISF_1$-neural (left) and $EEG_{resp}$-neural (right) coupling. Ordinary least squares (OLS) fits are marked by dashed lines. The histograms represents OLS residual distances and the bar-plots represents the autocorrelation function.

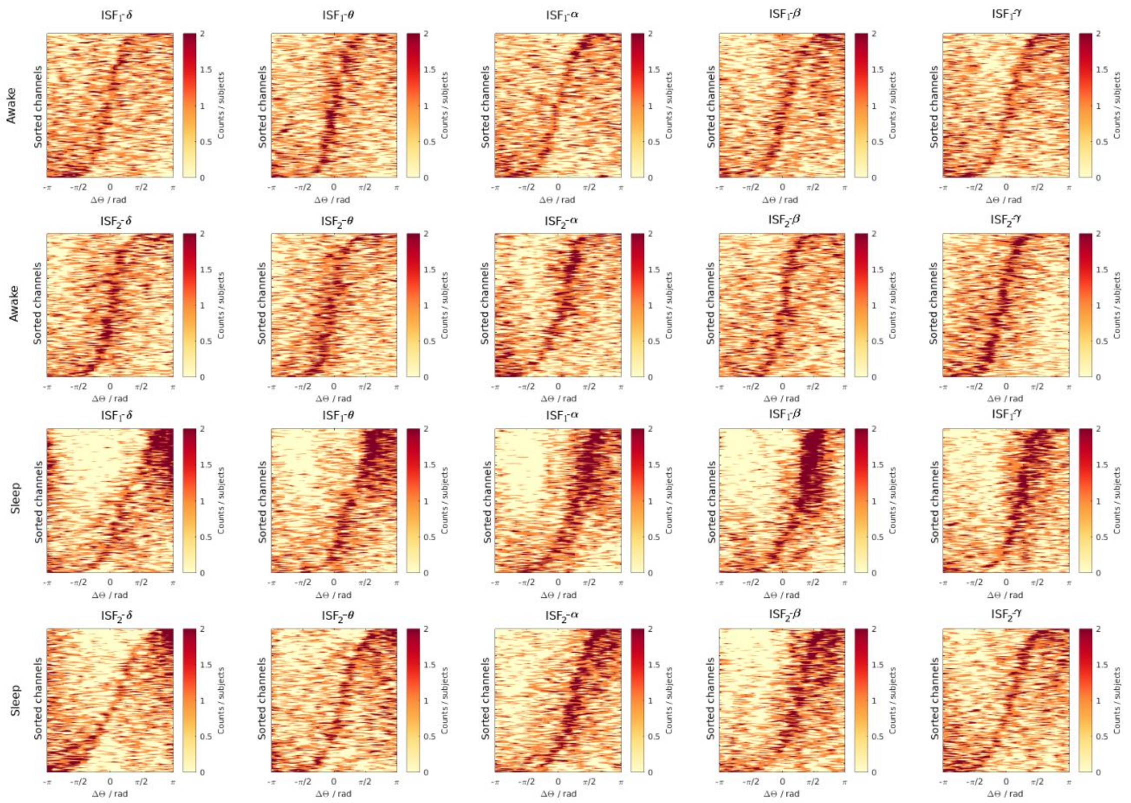


**Figure S6.** Probability estimates of the average phase differences between ISF phase and neural amplitudes. Channels on the y-axis are sorted to ascending order following the median phase difference. Greek letters indicate fast neural bands (delta, theta, alpha, beta, gamma).

### SI References


Ceguerra R v., Lizier JT, Zomaya AY (2011) Information storage and transfer in the synchronization process in locally-connected networks. In: 2011 IEEE Symposium on Artificial Life (ALIFE), pp 54–61. IEEE.

Lachaux J-P, Rodriguez E, Martinerie J, Varela FJ (1999) Measuring phase synchrony in brain signals. Hum Brain Mapp 8:194–208 Available at: https://onlinelibrary.wiley.com/doi/abs/10.1002/%28SICI%291097-0193%281999%298%3A4%3C194%3A%3AAID-HBM4%3E3.0.CO%3B2-C.

Lobier M, Siebenhühner F, Palva S, Palva JM (2013) Phase Transfer Entropy: A novel phase-based measure for directed connectivity in networks coupled by oscillatory interactions.
